# A *Drosophila* model of diabetic neuropathy reveals a crucial role of proteasome activity in the glia

**DOI:** 10.1101/2022.09.22.509008

**Authors:** Mari Suzuki, Hiroshi Kuromi, Mayumi Shindo, Nozomi Sakata, Naoko Niimi, Koji Fukui, Minoru Saitoe, Kazunori Sango

## Abstract

Diabetic peripheral neuropathy (DPN) is the most common chronic, progressive complication of diabetes mellitus. The main symptom is sensory loss; the molecular mechanisms are not fully understood. We found that *Drosophila* fed a high-sugar diet, which induces diabetes-like phenotypes, exhibit impairment of noxious heat avoidance. The impairment of heat avoidance was associated with shrinkage of the leg neurons expressing the *Drosophila* transient receptor potential channel Painless. Using a candidate genetic screening approach, we identified *proteasome modulator 9* as one of the modulators of impairment of heat avoidance. We further showed that proteasome inhibition in the glia reversed the impairment of noxious heat avoidance, and heat-shock proteins and exosome secretion in the glia mediated the effect of proteasome inhibition. Our results establish *Drosophila* as a useful system for exploring molecular mechanisms of diet-induced peripheral neuropathy and propose that the glial proteasome is one of the candidate therapeutic targets for DPN.

## INTRODUCTION

Diabetes mellitus (DM) is a public health problem that currently affects over 537 million people globally; the number is estimated to rise to approximately 643 million (11.3% of the population) by 2030 (International Diabetes Federation, 2021). Diabetic peripheral neuropathy (DPN) is the most common chronic and progressive complication of DM (Feldman et al., 2019; Jankovic et al., 2021). DPN is a major cause of chronic pain and paresthesia, but the most common symptom is a sensory loss that is a dominant risk factor for foot ulceration, leading to infections and toe or foot amputation. Despite intensive research on the effects of hyperglycemia on nerve function, the molecular mechanisms underlying DPN are still largely unknown, and effective therapies for DPN have not been established.

The progression of DPN involves damage to the peripheral sensory nervous system; thus, the vast majority of research on DPN has focused on neurons. However, it is noteworthy that peripheral sensory neurons are closely associated with glial cells; the axons are enclosed by Schwann cells and the neuronal soma is completely covered by satellite glial cells. Glial cells are indispensable for neurons, as they support the structure and function of neurons and promote neuronal survival. Recent work has suggested a role for glial cells in DPN. Patients with DPN show segmental demyelination without prominent axonal degeneration (Mizisin, 2014). Morphological changes, perturbations of metabolic pathways, such as the polyol pathway, and activation of an immune-like phenotype have been reported in Schwann cells of diabetic rodents (Goncalves et al., 2018). Satellite glial cells in dorsal root ganglia are also found to be activated in mouse and rat models (Hanani et al., 2014). These data suggest that glial cells may participate in the development of DPN. However, causation studies in humans are challenging to perform, and comprehensive genetic studies in mice are labor-intensive and time-consuming. Therefore, the causative role of glial cells and the basic molecular mechanisms that may underlie their contribution to DPN remain difficult to elucidate.

*Drosophila melanogaster* is a suitable animal model for genetic analyses and is also useful for the study of diabetes because the organs and molecular regulators of energy metabolism in *Drosophila* are mostly analogous to those in humans (Chatterjee and Perrimon, 2021). A high-sugar diet (HSD) has been used to study type 2 diabetes and its complications in *Drosophila*; like humans, adult flies fed the HSD develop hyperglycemia, insulin resistance, and obesity (Morris et al., 2012). The HSD also leads to heart and podocyte dysfunction, which resemble diabetic cardiomyopathy and diabetic nephropathy (Na et al., 2013, 2015). *Drosophila* has also been used to study the mechanisms of nociception, which allows animals to detect and avoid potentially harmful stimuli. Genes for mechanisms that regulate pain in mammals, such as transient receptor potential (TRP) channels and PIEZO channels, are conserved in *Drosophila* (He et al., 2022). In addition to the physiology of nociception, pathomechanisms of aberrant nociception have been studied in fly models. For example, nerve injury can lead to neuropathic pain by a loss of central inhibition in adult flies (Khuong et al., 2019). Fly larvae exhibit transient nociceptive sensitization following UV-induced injury, and HSD feeding or insulin receptor mutation interferes with recovery (Im et al., 2018). However, whether dietary-induced diabetic conditions induce sensory disturbances in adult flies remains unknown.

Here, we report that the HSD leads to DPN-like phenotypes in adult *Drosophila*. Flies reared on the HSD exhibited a reduction in the response behavior to noxious heat, which can be suppressed by treatment with antidiabetic drugs. In a candidate genetic screening approach, we identified the *Drosophila* homologue of *proteasome modulator 9* (*PSMD9*), one of the DPN-associated genes (Gragnoli, 2011a), as one of the modifier genes. We also found that proteasome inhibition in glial cells suppresses impairment of the noxious heat response. Moreover, heat-shock protein (HSP) DNAJ1 and exosome secretion in glial cells mediate the protective effect of proteasome inhibition. Our results suggest that glial proteasome activity is important for dietary sugar-induced sensory impairments. This new, genetically tractable model of DPN will be useful for understanding the molecular mechanisms of DPN.

## RESULTS

### Adult *Drosophila* fed an HSD exhibit a reduction in noxious heat avoidance behavior

To investigate whether high dietary carbohydrate induces sensory impairment in *Drosophila*, we fed adult wild-type (WT) flies a standard cornmeal-yeast-glucose diet (normal-sugar diet, NSD) or an HSD. The HSD contained 2.9-fold increased levels of total carbohydrates from the addition of 30% sucrose to the NSD (Table S1). We first examined the metabolic and systemic features of HSD-fed flies. These flies exhibited increased hemolymph sugar and decreased insulin sensitivity by 14 days (Figure S1A– D). The HSD induced an increase in *Insulin-like peptide 5* (*Ilp5*) expression on day 2, which returned to the basal level later (Figure S1E). Lipid droplets in the abdominal fat body were enlarged in the HSD-fed flies (Figure S1F), while total body weight was decreased (Figure S1G). HSD feeding significantly shortened the lifespan (Figure S1H). These results indicate that HSD feeding in adult flies induces DM-like metabolic features, as reported previously (Morris et al., 2012; Na et al., 2013; van Dam et al., 2020).

We then investigated whether flies fed the HSD exhibit sensory impairment. Avoidance behavior is dependent on nociception, the sensory process for detecting noxious or damaging stimuli. To evaluate thermosensory function, we developed a heat avoidance test, in which the heat band zone was set as the heat barrier to the flies climbing up by negative geotaxis (Figure 1A). The NSD-fed flies showed minimal avoidance responses when the heat band zone was set at 25°C, whereas they showed avoidance responses as the heat band zone temperature was increased to 38°C, 40°C, or 42°C) (Figure 1B). These heat avoidance features of NSD-fed flies were stably observed until at least 21 days after eclosion. HSD-fed flies also showed similar heat avoidance features until 7 days after feeding; however, the response to noxious heat (40°C and 42°C) was significantly decreased at 14 and 21 days (Figure 1B). Heat avoidance at 25°C and 38°C did not change over time, suggesting that HSD-induced avoidance impairment is specific to the response to noxious heat. In contrast to heat avoidance behavior, the locomotor function of flies fed the HSD was comparable to that of flies fed the NSD (Figure 1C). The effects of dietary sucrose on impairment of heat avoidance were dose-dependent and tended to be correlated with total hemolymph sugar concentration (Figure S2A–D). However, feeding flies the NSD without glucose (sugar-free), which leads to a decrease in hemolymph sugar, did not affect heat avoidance behavior (Figure S2A–D), suggesting that there is a threshold concentration of hemolymph sugar for inducing impairment of heat avoidance. Flies fed the natural banana diet with a carbohydrate content of approximately 22.5% (Table S1) showed similar heat avoidance profiles to flies fed the NSD (Figure S2E), suggesting that the heat avoidance profile of the NSD can be considered natural. Decreased heat avoidance behavior was also induced by other dietary saccharides, such as trehalose, glucose, and fructose (Figure 1D). HSD-induced impairment of heat avoidance was not observed when the antidiabetic drugs pioglitazone and metformin were administered at the same time as HSD feeding (Figure 1E). These results suggest that impairment of heat avoidance is caused by diabetes-like dysregulation of carbohydrate metabolism. Male WT flies had a lower basal avoidance rate than females, but impairment of heat avoidance was induced by the HSD (Figure 1B and Figure S3A). Unchanged locomotor function (Figure S3B) and decrease in body weight (Figure S3C) were similar to those in female flies (Figure 1C and Figure S1G), whereas the lifespan was not affected by the HSD in male flies (Figure S3D).

**Figure 1.**
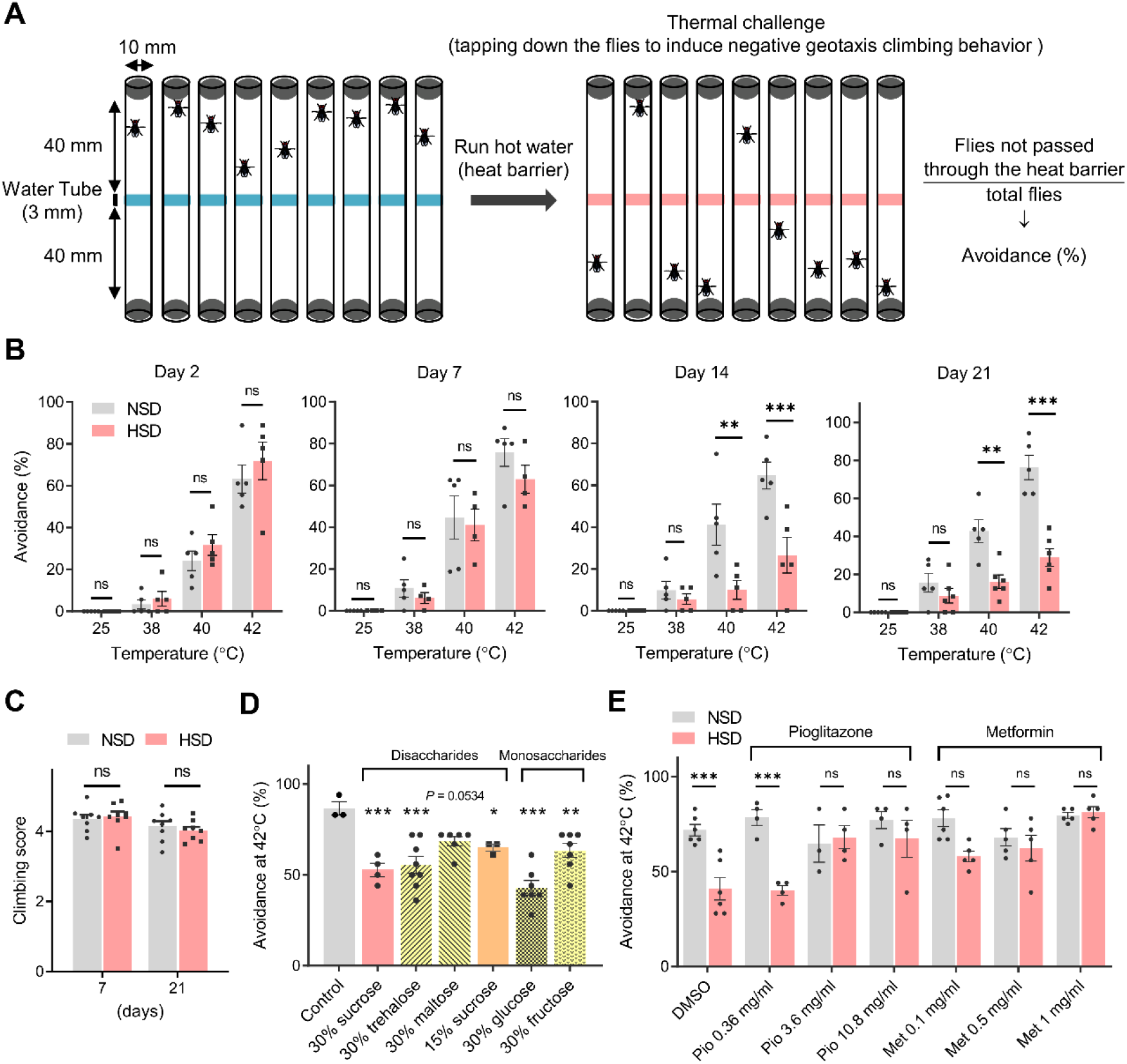
Adult *Drosophila* fed a high-sugar diet (HSD) exhibit a reduction in noxious heat avoidance behavior. (A) Schematic representation of the heat avoidance test in adult *Drosophila*. (B) Avoidance of noxious heat (40°C and 42°C), but not avoidance of subnoxious heat (25°C and 38°C), was significantly decreased in wild-type flies after HSD feeding for 14 and 21 days. NSD, normal-sugar diet. Two-way ANOVA post hoc Tukey’s test (N = 5–6, 7–9 flies per replicate). (C) Locomotor function assessed by the climbing assay was not affected by HSD. Two-way ANOVA post hoc Tukey’s test (N = 8, 10–20 flies per replicate). (D) Effects of dietary trehalose, maltose, glucose, and fructose on noxious heat avoidance. One-way ANOVA post hoc Dunnett’s test (each group was compared with the control, N = 3– 8, 7–9 flies per replicate). (E) HSD-induced impairment of noxious heat avoidance was suppressed by the antidiabetic drugs pioglitazone (Pio) and metformin (Met). Pioglitazone and metformin were administered at the same time as HSD feeding. Two-way ANOVA post hoc Tukey’s test (N = 3–6, 7–9 flies per replicate). Data are represented as means ± SEM. ns, not significant, \**P* < 0.05, \*\**P* < 0.01, \*\*\**P* < 0.001. DMSO, dimethyl sulfoxide.

### HSD-fed flies display shrinkage of leg sensory neurons

We next examined whether heat nociceptive neurons were affected by the HSD. Painless (Pain) is one of the TRP ion channel family required for the avoidance of noxious heat above 40°C in adult *Drosophila* (Khuong et al., 2019; Xu et al., 2006). Requirement of Pain for avoidance of noxious heat was confirmed in our test system, where a homozygous mutant of *pain* (*pain*^*EP*^) exhibited significant reduction of heat avoidance at 42°C (Figure 2A). Moreover, when we blocked synaptic output from Painless-expressing (*pain*+) neurons by expressing tetanus toxin (TNT) (Figure 2B) or temperature-sensitive shibire (shi^ts^) (Figure 2C) under control of *painless-GAL4* (*pain-GAL4*), the flies showed reduced heat avoidance even when fed the NSD. It has been reported that *pain*+ neurons are localized in the legs, thoracic ganglia (ventral nerve cord), and brain (Xu et al., 2006). Therefore, we next sought to examine whether HSD-induced impairment of the thermal response was due to dysfunction in peripheral regions other than the brain. To test this, we performed the hot plate test on decapitated flies. Decapitated NSD-fed WT flies showed jumping or tumbling in response to noxious heat (42ºC), as reported previously (Ohashi and Sakai, 2018), and the noxious heat response was significantly decreased in HSD-fed WT flies (Figure 2D). On the other hand, *pain*^*EP*^ mutant flies were less responsive to noxious heat, and the HSD did not further reduce the response (Figure 2D). These results suggest that disturbance of *pain*+ neurons in the peripheral regions is responsible for HSD-induced impairment of noxious heat avoidance.

**Figure 2.**
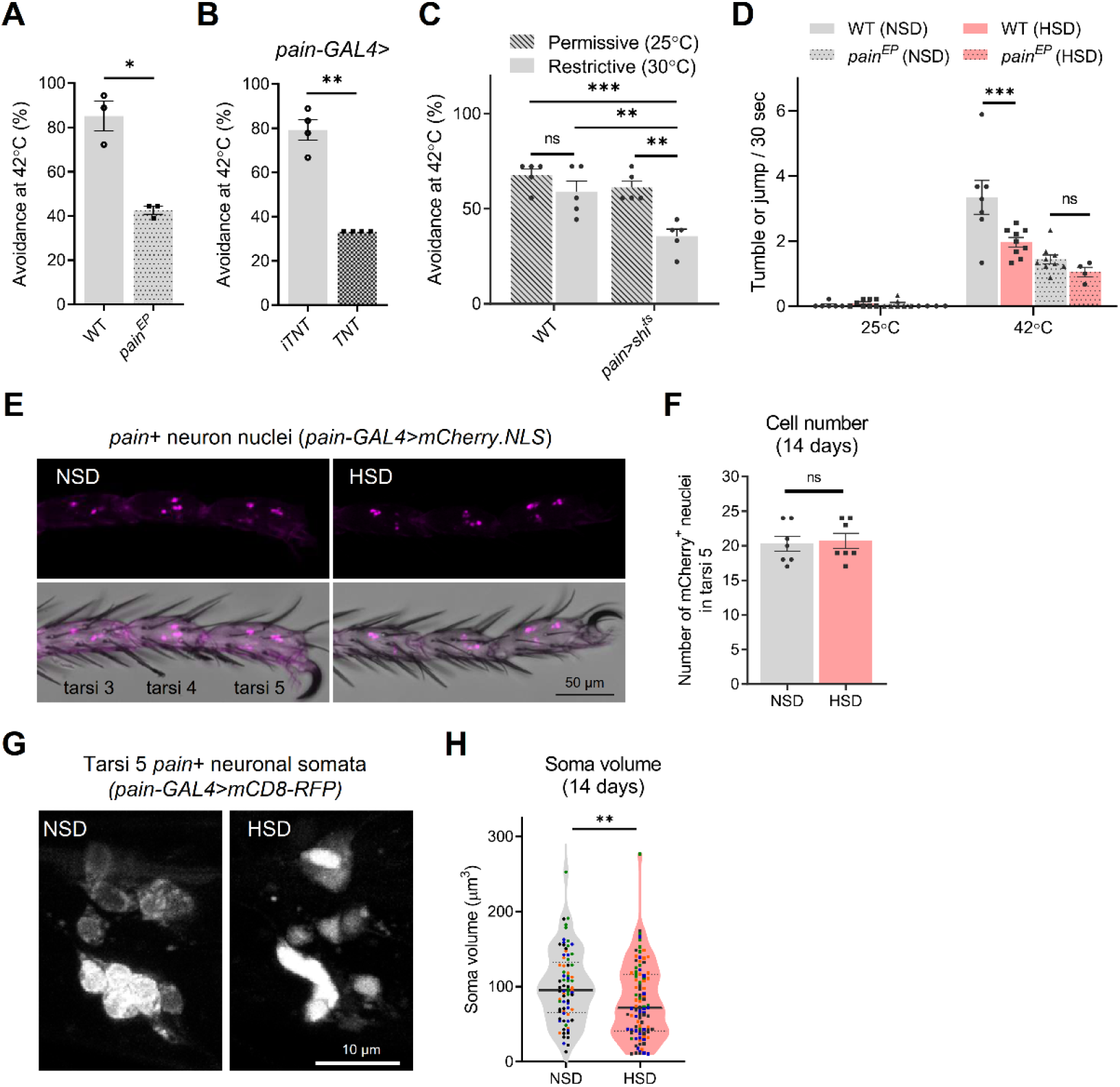
Adult *Drosophila* fed a high-sugar diet (HSD) exhibit a shrinkage of Painless-expressing leg sensory neurons. (A) The *Drosophila* transient receptor potential channel homologue Painless (Pain) is required for noxious heat avoidance. *Painless* mutant flies (*pain*^*EP*^) fed a normal-sugar diet (NSD) were tested. Welch’s *t* test (N = 3, 7–9 flies per replicate) WT, wild type. (B) Synaptic output from Pain-expressing (*pain*+) neurons is required for noxious heat avoidance. Flies expressing an active tetanus toxin (TNT) or an inactive tetanus toxin (iTNT) under control of the *painless* (*pain*)*-GAL4* fed NSD were tested. Welch’s *t* test (N = 4, 7–9 flies per replicate). (C) Expression of temperature-sensitive shibire^ts^ (shi^ts^) in *pain+* neurons significantly decreased heat avoidance at a restrictive temperature (30°C), but not a permissive temperature (25°C). Two-way ANOVA post hoc Tukey’s test (N = 5, 7–9 flies per replicate). (D) Hot plate test of decapitated flies. WT flies and *pain*^*EP*^ flies fed NSD or HSD were decapitated, and the noxious heat response was evaluated. Two-way ANOVA post hoc Tukey’s test (N = 4–9, 9 flies per replicate). (E) Nuclei of the leg *pain*+ neurons were visualized by expression of nuclear-localized mCherry (mCherry.NLS). (F) The cell number of the *pain*+ neurons in tarsi 5 was not changed by HSD feeding. Unpaired *t* test (N = 7 flies). (G) HSD results in shrinking of the *pain*+ neuron soma. Soma morphology of the *pain+* neurons was visualized by the expression of membrane-tethered red fluorescent protein (mCD8-RFP). (H) Soma volume of leg *pain*+ neurons was reduced in HSD-fed flies. Unpaired *t* test (N = 77 cells for NSD, 95 cells for HSD, 15–29 cells of four flies were quantified. Data points derived from each individual are distinguished by symbol color). Data are represented as means ± SEM. ns, not significant, \**P* < 0.05, \*\**P* < 0.01, \*\*\**P* < 0.001. All flies used were 2 weeks of age.

Therefore, we performed morphological analyses of *pain*+ neurons in the legs. Nuclei of the leg *pain*+ neurons were visualized by expression of nuclear-localized mCherry (mCherry.NLS) by *pain-GAL4*. We found 17–24 mCherry-positive nuclei in tarsi 5 of the T1 leg, and neuronal loss was not observed after 14 days of HSD feeding (Figure 2E and F). Morphological observation of neuronal cell bodies was performed by expressing membrane-tethered red fluorescent protein (mCD8-RFP) (Figure 2G). Although we could hardly observe projections of the *pain*+ neurons in the legs, we found that the neuronal somata became shrunken, with unclear boundaries and faint appearance, upon HSD feeding (Figure 2G and H). These results suggest that the impairment of heat avoidance in HSD-fed flies was caused not by neuronal death but by dysfunction of *pain*+ neurons.

### Proteasome inhibition in glial cells suppresses impairment of thermal nociception

To identify genetic modifiers of HSD-induced impairment of heat avoidance, we performed candidate screening for 13 genes (18 lines), which were selected on the basis of their functions in glucose metabolism (insulin-like signaling, glycolysis, polyol pathway, and hexosamine pathway), stress and cell death signaling, association with human DPN, and neurodegenerative diseases (Table S2). We used RNAi-mediated knockdown or overexpression flies under the control of the ubiquitously expressed driver *daughterless* (*da*)*-GAL4* or mutant flies to manipulate candidate gene expression systemically. Testing control flies, including WT, mCherry-RNAi, and EGFP overexpression flies, over several different days showed that the vast majority of NSD-fed control flies avoided the 42ºC noxious heat band, with a mean heat avoidance rate of 76.8% ± 9.2% SD, whereas HSD-fed control flies had a markedly reduced heat avoidance rate (35.3% ± 9.9% SD) (Figure S4A). On the basis of these data, we set our reference range with ± 2SD as 58.4%–95.1%. For this range, we consistently observed impairment of heat avoidance of the HSD-fed control flies (Figure S4A). We then performed the heat avoidance test with the test flies and identified six genes in seven fly lines in which the heat avoidance rates for both the NSD- and the HSD-fed flies are included in the reference range (Figure S4A and B).

From this screening, we picked out *CG9588* (*Drosophila proteasome modulator 9, dPSMD9*), an orthologue of human *proteasome modulator 9* (*PSMD9*). It has been suggested that polymorphisms of the *PSMD9* gene are associated with the risk of DPN (Gragnoli, 2011a; Salcini et al., 2021), but experimental validation and mechanism studies remain to be performed. We confirmed that *dPSMD9* knockdown suppressed impairment of heat avoidance in HSD-fed flies by using a second independent RNAi line (Figure 3A). These two RNAi lines resulted in approximately 41% and 58% reduction of whole body *dPSMD9* mRNA expression, respectively, when driven by the *da-GAL4* driver (Figure 3B). Because it has been reported that PSMD9 acts as a chaperone during the assembly of the 26S proteasome by binding with the Rpt4 and Rpt5 subunit (Kaneko et al., 2009; Saeki et al., 2009), we examined the effect of *Rpt4* and *Rpt5* knockdown. Similar to *dPSMD9* knockdown, knockdown of *Drosophila Rpt4* and *Rpt5* also diminished the effect of HSD (Figure 3C). On the basis of these results and a previous report that knockdown of *PSMD9* reduced the peptidase activity of the 26S proteasome in cultured cells (Kaneko et al., 2009), we hypothesized that proteasome activity has a key role in HSD-induced impairment of heat avoidance. To test this hypothesis, we investigated the effect of ixazomib, an oral proteasome inhibitor used for the treatment of multiple myeloma. Ixazomib treatment increased the heat avoidance of HSD-fed flies in a dose-dependent manner, and the reduction of HSD-induced heat avoidance completely disappeared at a dose of 10 μg/ml (Figure 3D). Furthermore, ixazomib treatment reversed the shrinkage of leg *pain*+ neurons (Figure 3E), suggesting that proteasome activity mediates the toxicity of HSD to *pain+* neurons. As well as chronic treatment, a shorter period of ixazomib pretreatment (days 0–7) and treatment after symptom onset (days 14– 21) also improved the reduction of heat avoidance (Figure 3F and G).

**Figure 3.**
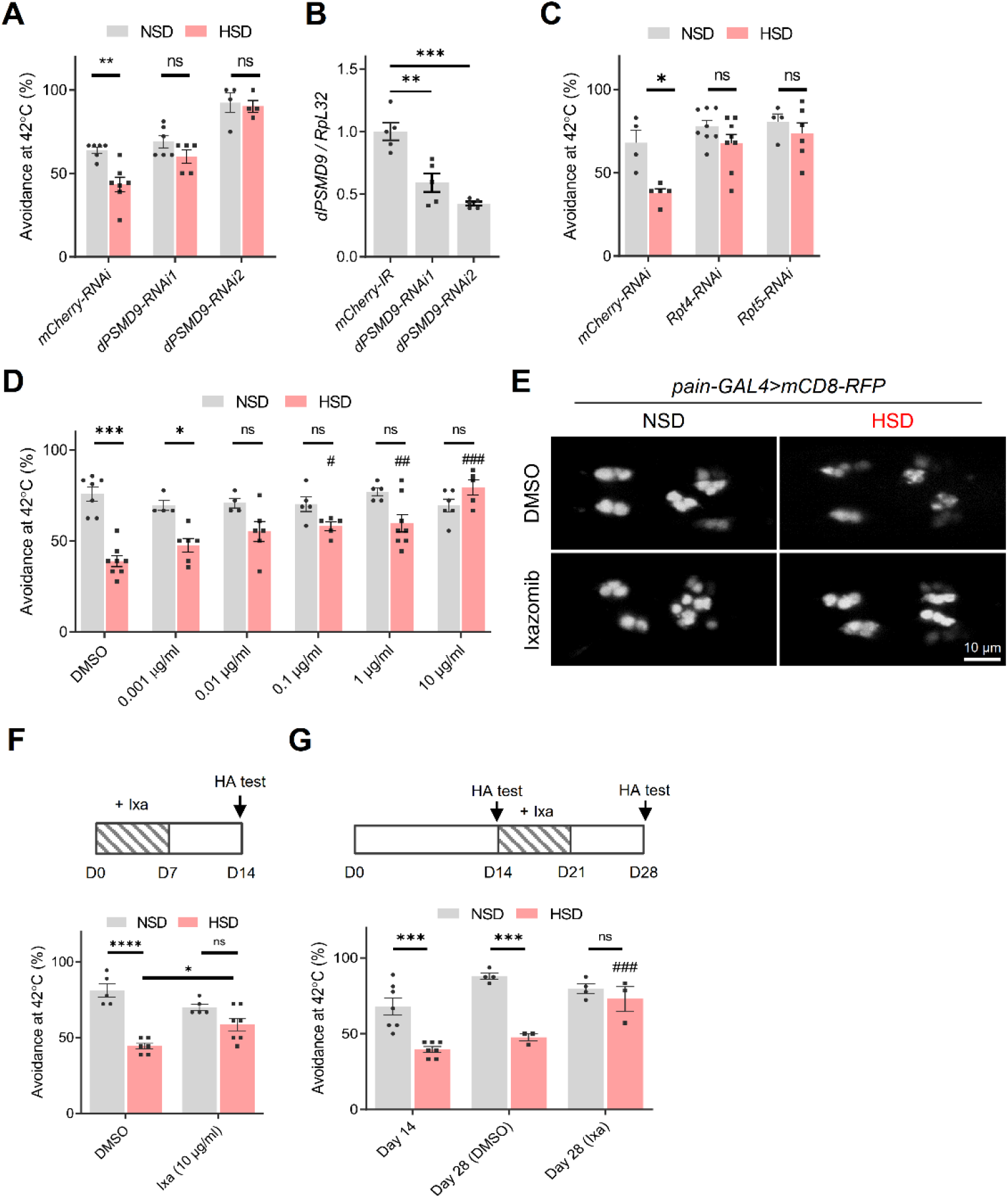
Knockdown of *Drosophila proteasome modulator 9* (*dPSMD9*) and proteasome inhibitor suppresses high-sugar diet (HSD)-induced impairment of heat avoidance. (A) Effects of *Drosophila proteasome modulator 9* (*dPSMD9*) knockdown on HSD-induced impairment of heat avoidance. Systemic RNAi-mediated knockdown was induced under control of the *daughterless* (*da*)*-GAL4*. Two-way ANOVA post hoc Tukey’s test (N = 4–7). NSD, normal-sugar diet. (B) Quantitative real-time polymerase chain reaction analysis of *dPSMD9* knockdown efficiency. RNAi-mediated knockdown was driven by the *da-GAL4* driver. One-way ANOVA post hoc Tukey’s test (N = 5 flies). (C) Effects of *Rpt4* and *Rpt5* proteasome subunit knockdown on impairment of heat avoidance. Two-way ANOVA post hoc Tukey’s test (N = 4–8). Knockdown was induced systemically under the control of *da-GAL4* (A–C). (D) The proteasome inhibitor ixazomib prevents HSD-induced impairment of heat avoidance of WT flies. Two-way ANOVA post hoc Tukey’s test (^##^*P* < 0.01, ^###^*P* < 0.001 vs. HSD with dimethyl sulfoxide (DMSO), N = 4–8). (E) HSD-induced shrinkage of the *pain*+ neurons was suppressed by ixazomib treatment (10 μg/ml). The images show mCD8-RFP signals in the tarsi 5 of fly legs. (F) Effects of ixazomib (Ixa) pretreatment (days 0–7). Two-way ANOVA post hoc Tukey’s test (N = 5–7). HA, heat avoidance. (G) Effects of ixazomib (Ixa) treatment after symptom onset (days 14–21). Two-way ANOVA post hoc Tukey’s test (N = 3–7). Data are represented as means ± SEM. HA, heat avoidance. ns, not significant, \**P* < 0.05, \*\**P* < 0.01, \*\*\**P* < 0.001.

We next investigated whether dPSMD9 and the proteasome act in neurons under HSD-induced impairment of heat avoidance. Unexpectedly, *dPSMD9* knockdown in *pain*+ neurons (*pain-GAL4*) as well as pan-neurons (*elav-GAL4*) did not suppress the impairment of heat avoidance (Figure 4A and B). In contrast, *dPSMD9* knockdown by glial-specific *repo-GAL4* abolished the effect of HSD (Figure 4C). Moreover, HSD-induced impairment of heat avoidance was suppressed by glia-specific proteasome inhibition, which was achieved by the expression of temperature-sensitive mutants of the 20S proteasome subunit β2 (Prosβ2^ts^) or both β2 and β6 (Prosβ2/6^ts^) at a restrictive temperature (Figure 4D). These results suggest that glial proteasome activity is responsible for HSD-induced impairment of heat avoidance.

**Figure 4.**
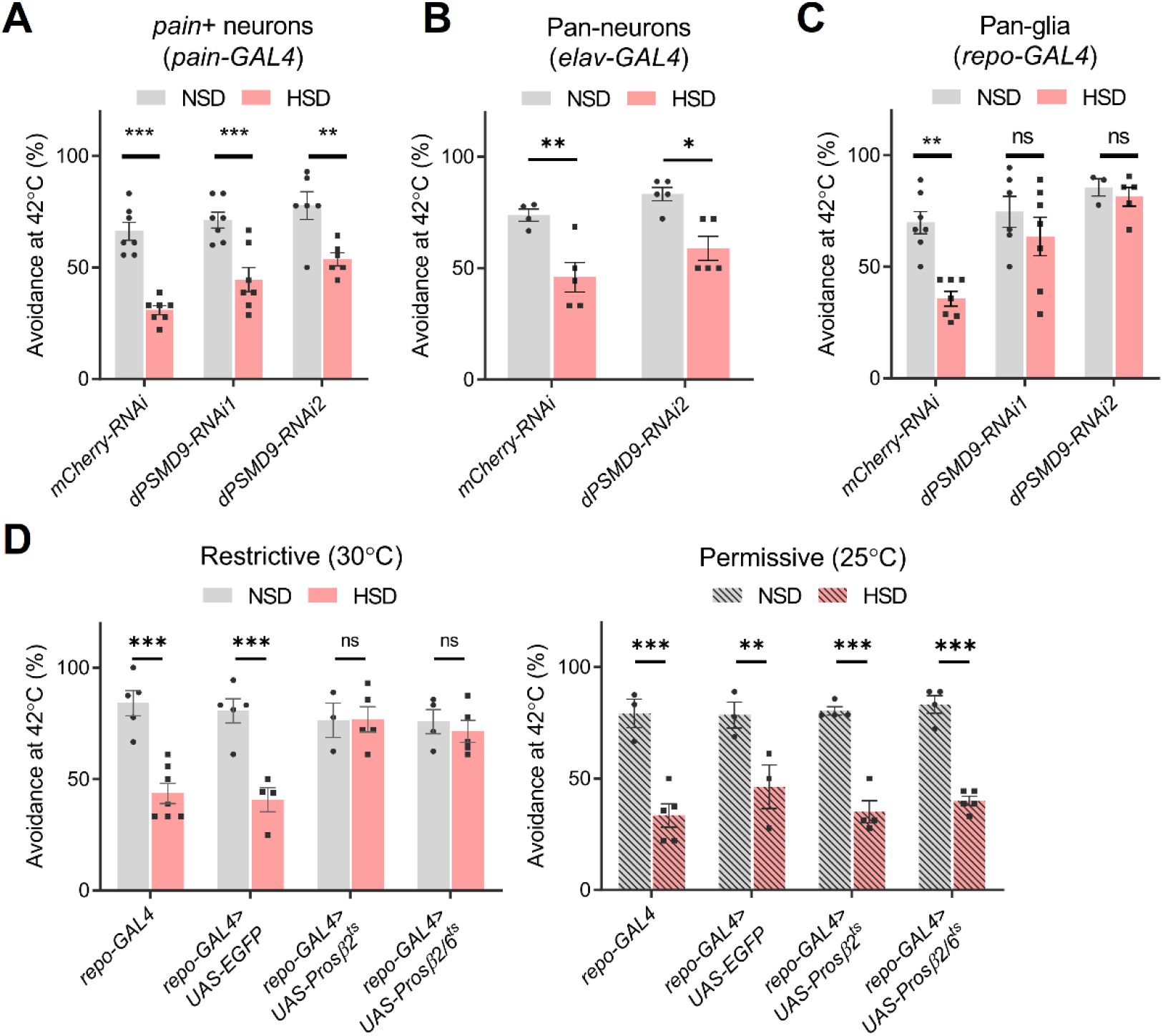
Proteasome inhibition in glial cells suppresses high-sugar diet (HSD)-induced impairment of heat avoidance. (A–C) Effects of tissue-specific *dPSMD9* knockdown on HSD-induced impairment of heat avoidance. *pain-GAL4* (A), *elav-GAL4* (B), and *repo-GAL4* (C) drivers were used for the knockdown in *pain*+ neurons, pan-neurons, and pan-glial cells, respectively. Two-way ANOVA post hoc Tukey’s test (N = 6–7 for A, N = 4–5 for B, N = 3–7 for C). NSD, normal-sugar diet. (D) HSD-induced impairment of heat avoidance is alleviated by glia-specific proteasome inhibition. Flies expressing the dominant temperature-sensitive mutants of the 20S proteasome subunit, β2 (*repo-GAL4>UAS-Prosβ2*^*ts*^), or both β2 and β6 (*repo-GAL4>UAS-Prosβ2/6*^*ts*^), in glial cells were used for the heat avoidance test at either a restrictive (30°C) or a permissive (25°C) temperature. Two-way ANOVA post hoc Tukey’s test (N = 3–7). Data are represented as means ± SEM. ns, not significant, \**P* < 0.05, \*\**P* < 0.01, \*\*\**P* < 0.001.

### Glial HSPs and exosome secretion mediate the protective effect of proteasome inhibition

To investigate how proteasome inhibition in glial cells suppresses impairment of heat avoidance, we employed isobaric tags for relative and absolute quantitation-mass spectrometry (iTRAQ-MS) analysis using cultured mouse Schwann cells. We identified 2190 proteins, and a total of 66 differentially expressed proteins were identified with false discovery rate (FDR)-corrected *P*-values < 0.05, including 23 down-regulated and 43 up-regulated in ixazomib-treated cells (Figure 5A, Table S3A and S3B). Metascape enrichment analysis revealed that the differentially expressed proteins were enriched for the pathway related to protein folding (Figure 5B, Table S3C). Correspondingly, HSPs such as HSP40, HSP70, heat-shock cognate 71 kDa protein (Hsc70), HSP90α, and HSP90β were markedly up-regulated (Figure 5C). Up-regulation of the polyubiquitinated proteins, HSP40, HSP70, and HSP90 was confirmed by immunoblotting analyses, while the amount of PSMD9 was not changed (Figure 5D and E).

**Figure 5.**
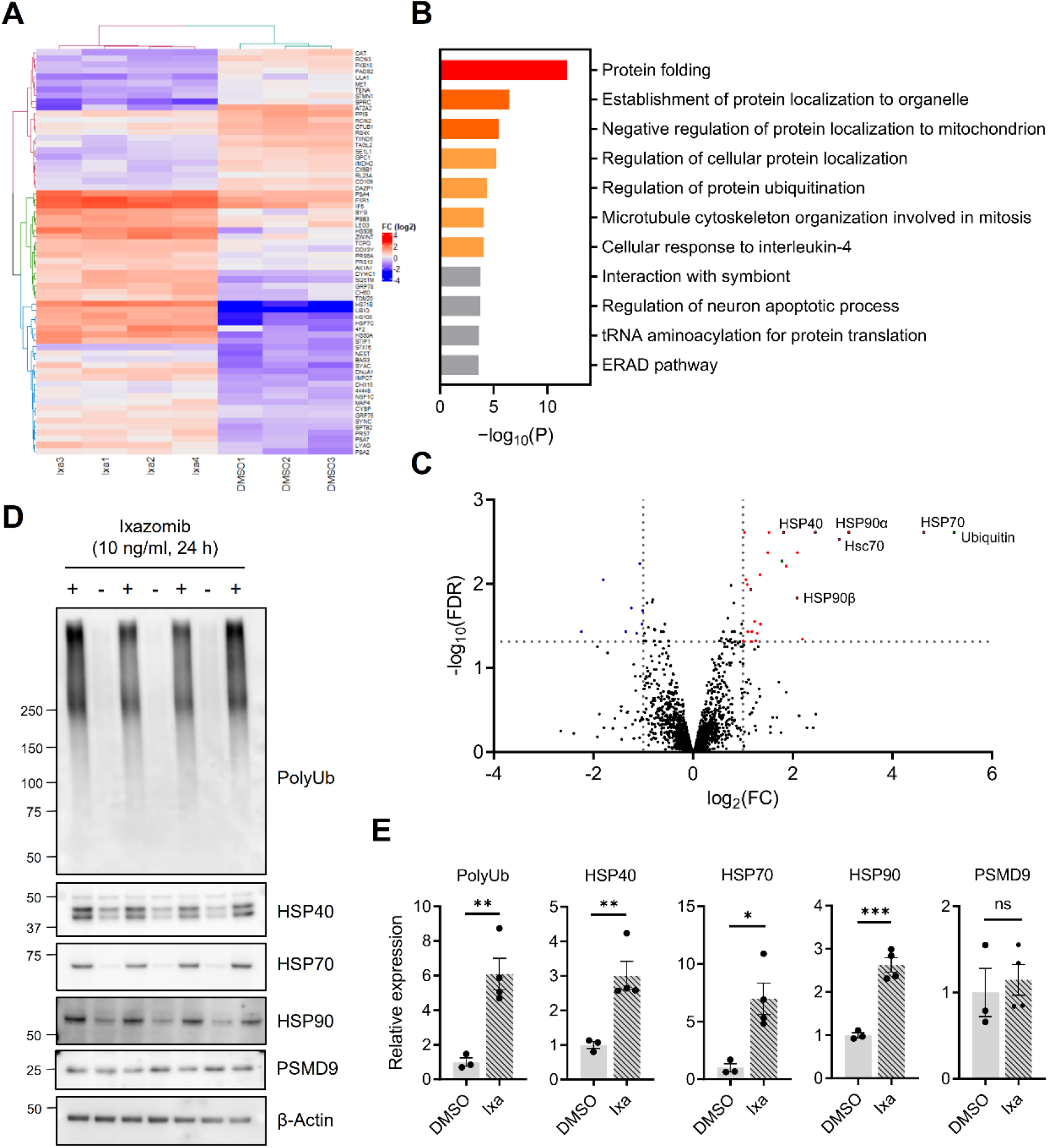
Ixazomib increases heat-shock proteins (HSPs) in mouse Schwann cells. Immortalized mouse Schwann cells treated with either 10 μg/ml ixazomib (Ixa) or dimethyl sulfoxide (DMSO) for 24 h were subjected to isobaric tags for relative and absolute quantitation-mass spectrometry (iTRAQ-MS) proteomics. (A) Overview of the 66 significantly changed proteins (false discovery rate [FDR]-corrected *P* < 0.05) in four ixazomib Ixa groups as compared with three DMSO groups. The heatmap represents the log_2_ (fold change [FC]) of the differentially expressed proteins in each group. The color key from blue to red represents the log_2_ (FC) from low to high. See Table S3A and B for details. (B) Enrichment analysis of differentially expressed proteins. The vertical axis represents the pathway category and the horizontal axis represents the enrichment score [–log_10_ (P)] of the pathway. See Table S3C for details. (C) Volcano plot representing log_10_ (FDR) as a function of log_2_ (FC). The proteins identified with an FDR-value < 0.05 and > 1.0-fold are highlighted in red (up-regulated) or blue (down-regulated), with the dotted lines representing the boundary for identification. (D) Immunoblotting analyses using antibodies against poly-ubiquitin (polyUb), HSP40, HSP70, HSP90, and proteasome modulator 9 (PSMD9). (E) Quantification of relative protein amounts in (D). β-Actin was used for internal control. Unpaired *t* test. Data are represented as means ± SEM. ns, not significant, \**P* < 0.05, \*\**P* < 0.01, N = 3 for DMSO, and 4 for ixazomib.

We then examined whether glial HSPs are required for the effect of ixazomib in *Drosophila*. We first investigated the effect of glia-specific knockdown of heat-shock transcription factor (Hsf), a master regulator of the heat-shock genes, including DNAJ1, Hsp70, and Hsp83 (Gonsalves et al., 2011). Hsf-RNAi flies showed a slight reduction in heat avoidance, even in the NSD condition, and the effect of ixazomib, which was seen in the control knockdown (mCherry-RNAi) flies, was abolished (Figure 6A). We also selected DNAJ1, one of the HSP40 family members, as a target for a knockdown experiment, because we had found a reduced expression level of *DNAJ1* in HSD-fed flies (Figure S5). In addition to the Hsf-RNAi flies, the effect of ixazomib was abolished in all three DNAJ1-RNAi lines (Figure 6A), suggesting the requirement of glial DNAJ1 for the effect of ixazomib. Interestingly, DNAJ1-RNAi2 and DNAJ1-RNAi3 themselves reduced heat avoidance, even in the NSD condition, and the reduction was not reversed by ixazomib (Figure 6A). In contrast, glial overexpression of DNAJ1 reversed the reduction in heat avoidance induced by HSD, suggesting that an increased amount of DNAJ1 in the glia is sufficient for the improvement of HSD-induced impairment of heat avoidance (Figure 6B).

**Figure 6.**
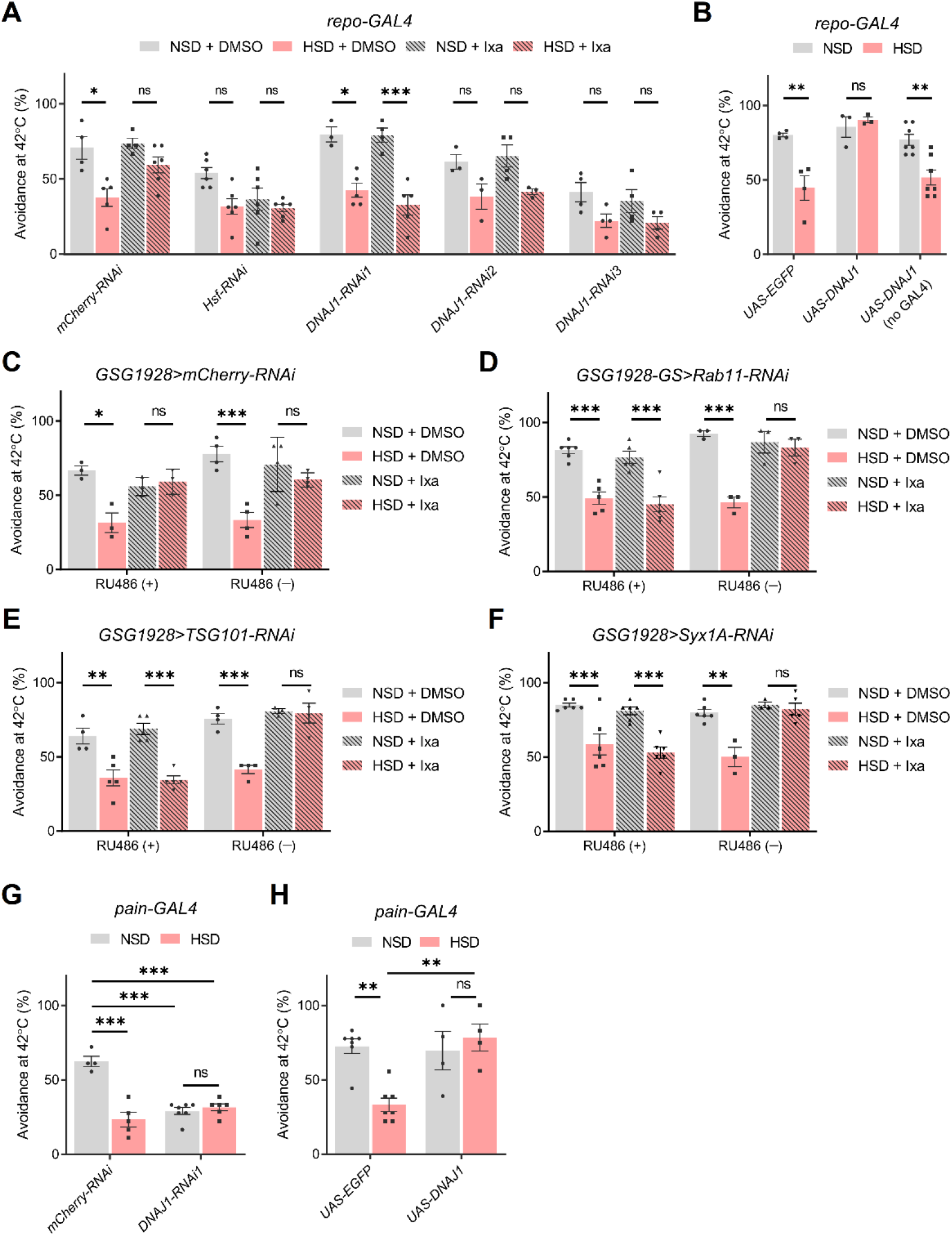
Glial heat-shock proteins (HSPs) and exosome secretion mediate the effect of ixazomib on high-sugar diet (HSD)-induced impairment of heat avoidance. (A) Knockdown of *DNAJ1* and *Heat-shock factor* (*Hsf*) in the glia abolished the effect of ixazomib (Ixa). Two-way ANOVA post hoc Tukey’s test (N = 3–6). DMSO, dimethyl sulfoxide; NSD, normal-sugar diet. (B) Glia-specific overexpression of DNAJ1 reversed the impairment of heat avoidance induced by HSD. Two-way ANOVA post hoc Tukey’s test (N = 3–7). (C–F) Glia-specific knockdown of the genes required for exosome release abolished the effect of ixazomib. Conditional knockdown of *Syx1A, Rab11*, and *TSG101* was induced by administration of RU486 after eclosion. Two-way ANOVA post hoc Tukey’s test (N = 3–4 for C, N = 3–6 for D, N = 3–5 for E, N = 3–6 for F). (G, H) The effect of *pain+* neuron-specific knockdown (G) or overexpression (H) of DNAJ1. Two-way ANOVA post hoc Tukey’s test (N = 4–6 for G, N = 4–7 for H). Data are represented as means ± SEM. ns, not significant, \**P* < 0.05, \*\**P* < 0.01, \*\*\**P* < 0.001.

We next investigated how the HSPs in glial cells rescue neurons. We previously demonstrated that HSPs are secreted via exosomes and contribute to the maintenance of protein homeostasis in different cells (Takeuchi et al., 2015). Thus, we hypothesized that exosomal transmission of HSPs contributes to the effect of ixazomib. To test this hypothesis, we focused on the genes related to exosomal release. *Rab11, Syntaxin 1A* (*Syx1A*), and *Tumor susceptibility gene 101* (*TSG101*) are known to function in multivesicular body transportation, membrane fusion with the plasma membrane, and the endocytosis sorting complex required for transport (ESCRT)-I, respectively (Tsai et al., 2019). Because *repo-GAL4*-driven knockdown of these genes resulted in lethality, we used the Gene-Switch system, in which transcriptional activity within the target tissues depends on the presence of the activator RU486 (Osterwalder et al., 2001). The *GSG1928* driver is reported to induce glial expression in the presence of RU486 (Nicholson et al., 2008); when we used this driver for conditional knockdown of *Rab11, TSG101*, and *Syx1A*, the effect of ixazomib was abolished only in the presence of RU486 (Figure 6C– F). These results suggest that exosomal secretion from the glia is required for the effect of ixazomib.

We further examined the role of DNAJ1 in *pain*+ neurons for noxious heat avoidance. DNAJ1 knockdown in *pain+* neurons reduced the heat avoidance even under NSD feeding, and HSD did not further reduce the heat avoidance (Figure 6G). On the other hand, overexpression of DNAJ1 in *pain+* neurons suppressed HSD-induced impairment of heat avoidance, but did not affect the heat avoidance under NSD feeding (Figure 6H). Collectively, these results support our hypothesis that glial cells supply DNAJ1, which is required for the proper function of the *pain+* neurons in thermal nociception, via exosomal secretion upon proteasome inhibition.

## DISCUSSION

This study shows that feeding of HDS induces DPN-like phenotypes in adult flies. We identified dPSMD9 as a modulator of HSD-induced impairment of nociceptive heat avoidance, and further found that inhibition of proteasome activity in glial cells reverses the impairment through HSPs and exosome secretion. This study provides a novel genetically tractable model for studying the molecular mechanisms of DPN and further highlights the role of glial proteasome activity under diabetic conditions.

Sensory disturbance of DPN is characterized by both positive symptoms, such as pain, and negative symptoms, such as loss of sensation. Although sensory loss is a common symptom, approximately 15%–25% of people with DM present with neuropathic pain (Shillo et al., 2019). One puzzling difference between our adult fly model and the previously reported larval fly model (Im *et al*., 2018) is the direction of sensory abnormalities. Although it has been suggested that positive symptoms change to negative symptoms as the disease progresses (Yagihashi et al., 2007), we did not observe significant thermal sensitization, even at early time points (Figure 1B). In fly larvae, HSD induces the persistence of thermal hypersensitivity but does not change the baseline thermal nociception and acute thermal hyperalgesia response after injury (Im et al., 2018), suggesting that the recovery process may be a key for induction of the sensitization. Because neuropathic sensitization can be induced by nerve injury in adult flies (Khuong et al., 2019), it will be of interest to determine whether HSD exacerbates or alleviates the sensitization of the nerve injury model. It has been reported that the majority of patients with painful DPN show sensory loss on clinical examination (Themistocleous et al., 2016), indicating that painful and painless DPN are not mutually exclusive. Therefore, establishing a painless DPN model in adult flies, in addition to the larval painful DPN model, would be important for further understanding of the molecular mechanisms of DPN.

Another phenotypic feature of our model is the sensory-dominant manifestation, which agrees well with the human DPN (Sloan et al., 2021). Sensory disturbance appeared, but motor disturbance did not occur until at least 3 weeks (Figure 1B and C), indicating a difference in vulnerability to HSD. This difference in vulnerability may be explained by the location of the sensory and motor neurons; in *Drosophila*, the cell bodies of sensory neurons are located in the periphery, as in mammals, and the cell bodies of motor neurons are located inside the ventral nerve cord, which is equivalent to the vertebrate spinal cord (Sánchez-Soriano et al., 2007). Because the spinal cord has a blood–spinal cord barrier, a physical barrier between the blood and spinal cord parenchyma, the neurons inside the spinal cord are protected against chemical imbalance. Although, at least to our knowledge, there is no clear evidence that the ventral nerve cord of the adult fly has an equivalent function to the blood–spinal cord barrier, the surface glia, which forms the blood–brain barrier in the *Drosophila* brain, is found to exist in the ventral nerve cord (Allen et al., 2020) and may provide a protective environment for the cells inside it.

The advantage of the fly DPN model is its application in genetic studies. For candidate genetic screening, we selected *Aldo-keto reductase 1B* (*Akr1B*), *Poly-(ADP-ribose) polymerase* (*Parp*), *Pelle* (*pll*), Relish (*Rel*), and *dPSMD9*, which are the *Drosophila* homologues of human DPN-associated genes or their signaling cascade molecules (asterisks in Table S2) (Gragnoli, 2011a; Gupta and Singh, 2017; Heesom et al., 1998; Nikitin et al., 2008; Rudofsky et al., 2004; Salcini et al., 2021; Sivenius et al., 2004). The genes other than *Parp* were identified as modulators according to our criteria (Figure S4). Aldose reductase (AR), one of the predicted homologues of Akr1B, is a late-limiting enzyme in the polyol pathway that converts glucose to sorbitol and fructose (Niimi et al., 2021). The pathological role of AR has been demonstrated in AR knockout mice (Ho et al., 2006), and an AR inhibitor epalrestat has been approved for DPN treatment in Japan (Hotta et al., 2012). Toll-like receptor 4 (TLR4) is another DPN-associated gene, and we have chosen downstream effectors *pll* and *Rel*, which are homologs of human *interleukin-1 receptor-associated kinase 4* (*IRAK4*) and *nuclear factor kappa B* (*NFkB*), respectively, because there is no definite fly homologue of TLR4. Although *pll* and *Rel* function in the independent immune signaling cascades, the toll signaling pathway and the immune deficiency signaling pathway, both genes are identified as modulators, suggesting the importance of immune signaling. Our finding that several human DPN-associated genes modulate HSD-induced impairment of heat avoidance further supports the validity of our fly DPN model.

A striking finding of our study is that inhibition of proteasome activity in the glia can alleviate HSD-induced neuropathy. Changes in proteasome activity under DM conditions vary among the reports. For instance, it has been reported that hyperglycemia impairs proteasome function by glycation with methylglyoxal in the kidney and aorta of type 1 DM mice (Queisser et al., 2010). Beta cells of patients with type 2 DM showed down-regulation of proteasome genes with ubiquitin accumulation and reduced proteasome activity (Bugliani et al., 2013). In contrast, there are reports that high glucose levels lead to an acceleration of proteasomal proteolysis in muscle, endothelial, renal, and retinal cells (Fernandes et al., 2004; Liu et al., 2012; Wang et al., 2006). It is interesting to note that administration of proteasome inhibitors, such as MG132 or PR-11, reversed alterations of endothelium-dependent vessel relaxation and NFκB–mediated proinflammatory response in the aorta, kidney, and retina of diabetic mice (Kong et al., 2017; Liu *et al*., 2012; Xu et al., 2007, 2012). However, changes in proteasome activity in the nervous system of diabetic animals have not been reported to date. Further study is required to determine whether diabetes enhances proteasome activity in the peripheral nervous system, including in a *Drosophila* model.

Current treatments for DPN focus on the control of blood glucose. Ixazomib is an FDA-approved orally available drug for multiple myeloma, but it is known to result in peripheral neuropathy as one of the adverse effects (Muz et al., 2016). Therefore, ixazomib would not be applicable to treat DPN, at least with the regimens approved for multiple myeloma. Although ixazomib at the concentrations we used did not affect basal noxious heat avoidance behavior (Figure 3D), it has been reported that knockdown of the proteasome alpha subunits PSMA in sensory neurons results in insensitivity to thermal nociception in fly larvae (Honjo et al., 2016). Moreover, chronic treatment with ixazomib at a dose of 10 μg/ml shortened the lifespan of flies both receiving the NSD and receiving the HSD (data not shown), a result similar to the previous report (Suraweera et al., 2012), implying that ixazomib is not suitable for long-term use. Therefore, seeking an optimized regimen or alternative interventions by elucidating the mechanism of the action of proteasome inhibition may be required.

We showed that the molecular chaperones are among the downstream effectors of the action of ixazomib. Although induction of molecular chaperones for DPN treatment has received some attention by other groups (Dobrowsky, 2016; Henstridge et al., 2014), it is a novel finding that induction of molecular chaperones in glial cells alone is sufficient for symptomatic treatment (Figure 6B). In addition, we showed that exosomal secretion of glial cells is also required (Figure 6D–F). Taken together with our data and a previous report of exosomal chaperone transmission (Takeuchi et al., 2015), we consider that proteasome inhibition induces exosomal transmission of HSPs from glial cells to sensory neurons, providing protection against HSD. However, we cannot rule out the possibility that other components of the exosome play a key role. It has been demonstrated that exosomal miRNA derived from glial cells regulates the growth of synaptic boutons of motor neurons and tracheal branches (Tsai et al., 2019), suggesting the possibility that the effect of exosomal secretion is mediated by other cells or tissues. It is interesting to note that the administration of Schwann cell-derived exosomes ameliorates peripheral neuropathy in diabetic mice (Wang et al., 2020). Since the glial cells also support neurons via lactate and alanine transport (Volkenhoff et al., 2015), detailed studies will be required to determine whether proteasomes, HSPs, and exosomes influence the glial metabolic support for neurons.

Although we assume that reduction of proteasome activity resulting from dPSMD9 knockdown is important for HSD-induced impairment of heat avoidance, it is also possible that loss of other dPSMD9 functions contributes to the phenotypes. One of the single nucleotide polymorphisms (SNPs) associated with DPN is E197G, which lies in close proximity to the PDZ domain (pt.108–195) (Gragnoli, 2011a). The PDZ domain is known as a protein–protein interaction module, and is reported to be required for interaction of PSMD9 with basic helix-loop-helix transcription factors E12 and E47 (Thomas et al., 1999) and inhibitor of nuclear factor κBα (IκBα) (Sahu et al., 2014), implying roles in insulin gene transcription and inflammation, respectively. The same SNPs are also reported to be associated with the risk of type 2 DM (Gragnoli, 2010; Gragnoli and Cronsell, 2007), diabetic nephropathy (Gragnoli, 2011b), anxiety (Gragnoli, 2014), and insomnia (Hao et al., 2015), implying diverse roles. In addition, trans-omics analyses of 107 genetically distinct mouse strains identified PSMD9 as a lipid regulatory protein in the plasma and liver (Parker et al., 2019). The authors showed that PSMD9 silencing led to a significant reduction in the synthesis of fatty acids in the liver. It will be interesting to see if this function is similar in glial cells and if it is similar through regulation of proteasome activity.

This study has shown that the regulation of proteasome and molecular chaperones in glial cells plays a crucial role in maintaining the function of sensory neurons under diabetic conditions. Although it remains to be determined which subtype of glial cells is important, the proteostasis mechanisms in glial cells can be targeted for treatment of DPN. With the development of genetic analysis technology, genetic susceptibility factors for DM and DPN are being identified, but the biological significance of these genetic factors has not yet been studied. Considering the convenience of the fly for creating libraries of multiple genetic models, *Drosophila* genetics coupled with mammalian validation will be a powerful system to investigate the molecular mechanisms leading to DPN.

### Limitations of the study

There are two major limitations of this study that should be addressed in future. First, although *Drosophila* provides a powerful model system for understanding the molecular mechanisms underlying DPN, the relevance and generalizability of these mechanisms to human disease remains to be demonstrated in mammalian experimental models. Second, this study lacks neurophysiological demonstration of dysfunction in the *pain*^+^ neurons.

## ACKNOWLEDGMENTS

We are grateful to the Bloomington *Drosophila* Stock Center and the Vienna *Drosophila* Resource Center. We thank Kumi Sumida for technical assistance. This work was supported in part by a Grant-in-Aid for Scientific Research (C) (19K07987 to M. S.) from the Ministry of Education, Culture, Sports, Science, and Technology, Japan; research grants from the Takeda Science Foundation (to M. S.); research grants from the Mochida Memorial Foundation for Medical and Pharmaceutical Research (to M. S.); and research grants from the Suzuken Memorial Foundation (to M. S.).

## AUTHOR CONTRIBUTIONS

M. Suzuki: conceptualization, data curation, funding acquisition, investigation, writing (original draft), writing (review and editing); H. K.: methodology; M. Shindo: data curation; N. S.: data curation, investigation; N. N.: resources; K. F.: supervision; M. Saitoe: supervision; K. S.: funding acquisition, project administration, supervision, writing (review and editing).

## STAR METHODS

### KEY RESOURCES TABLE

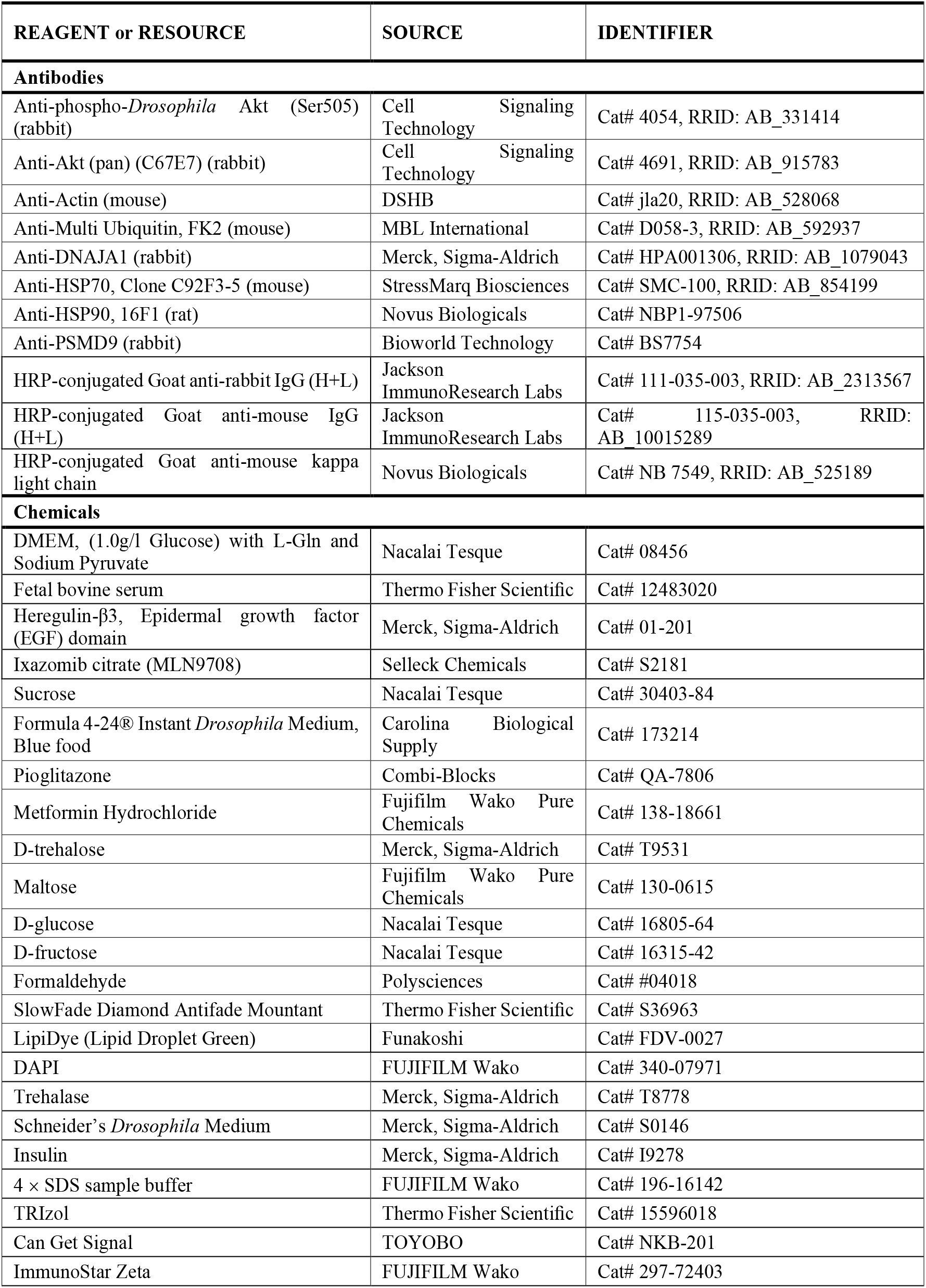
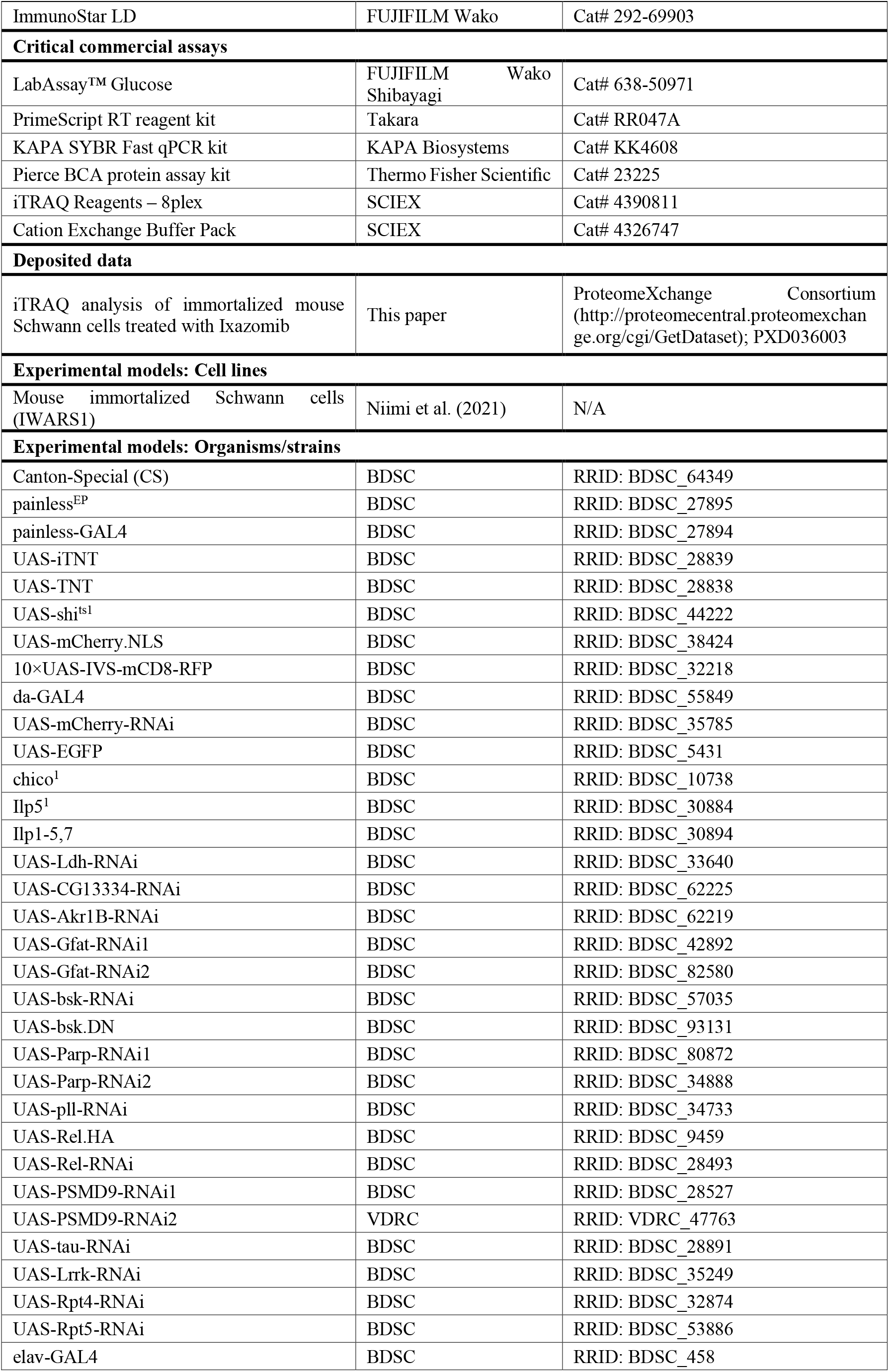
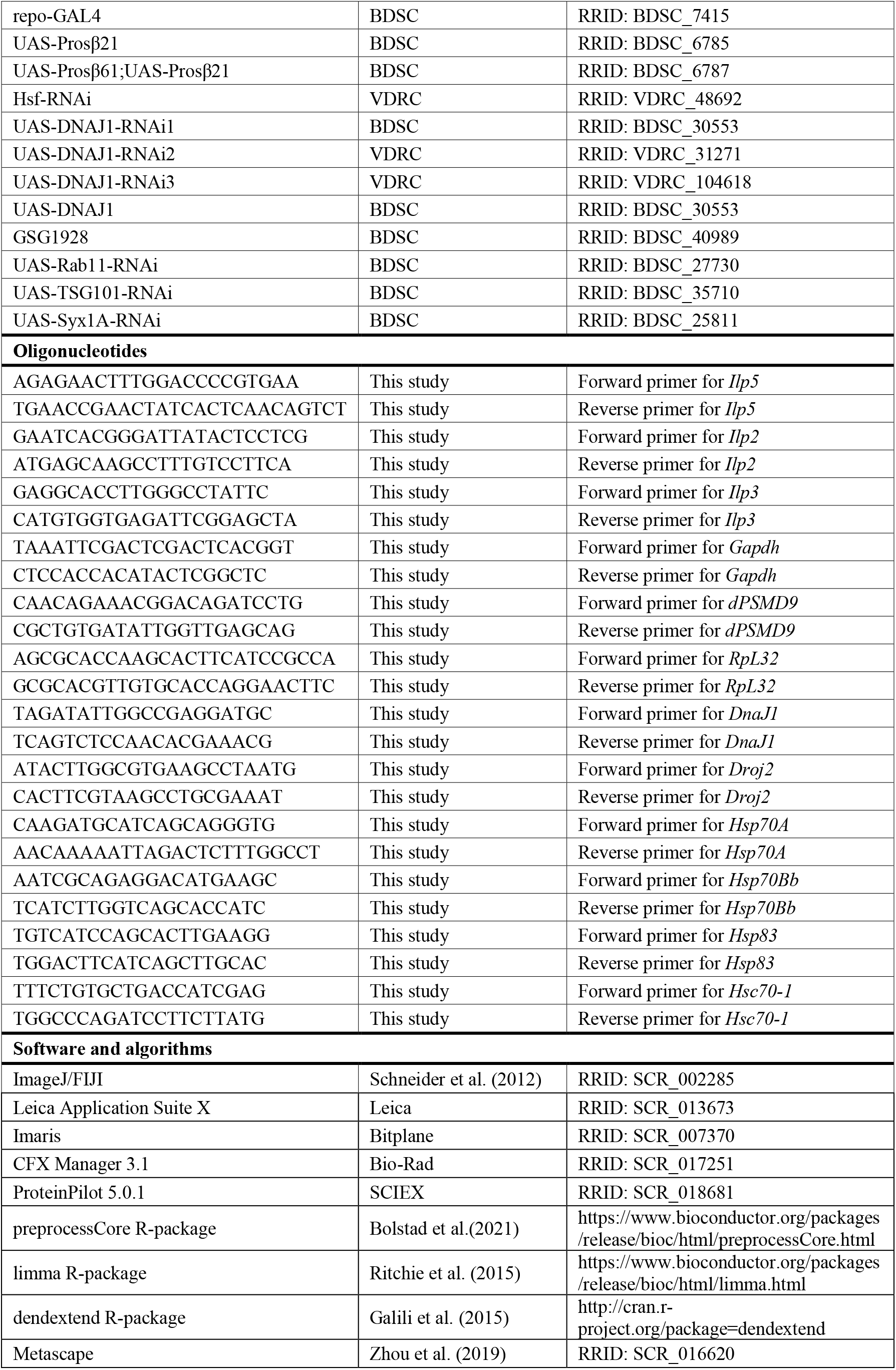
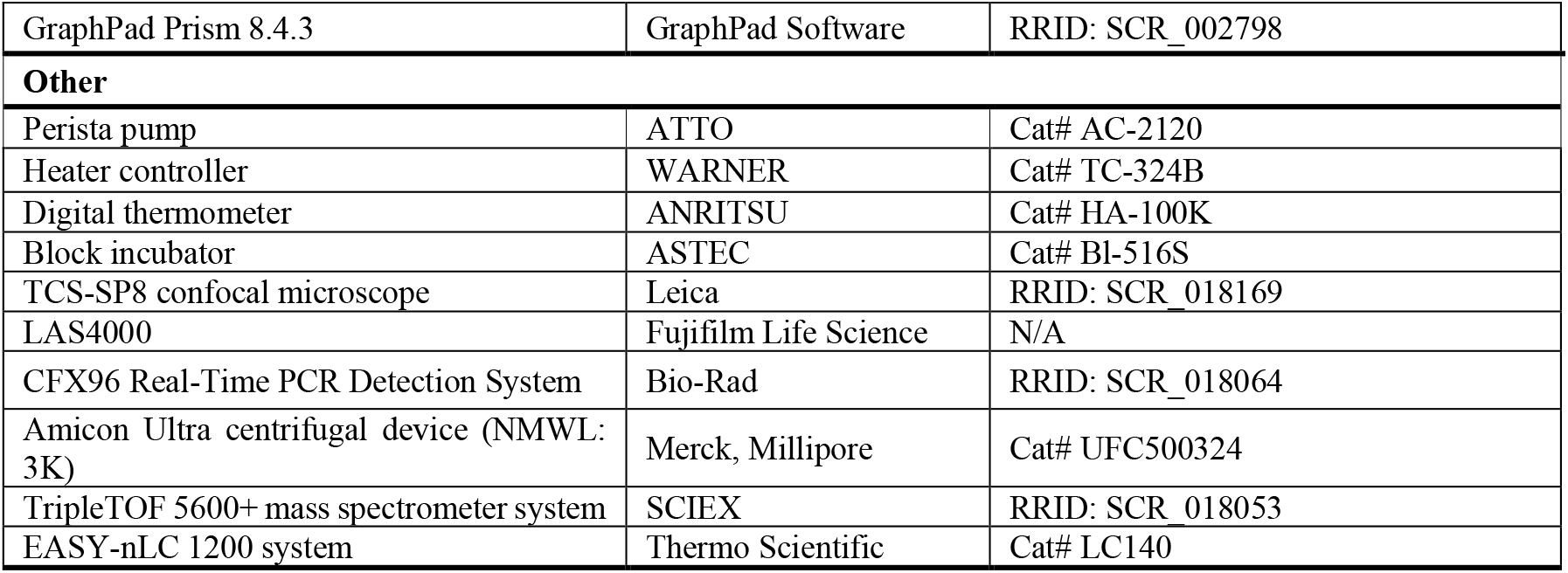

### RESOURCE AVALABILITY

#### Lead contact

Requests for further information, reagents, and resources should be directed to and will be fulfilled by the Lead Contact, Mari Suzuki.

#### Materials availability

Adult WT C57/BL6J mouse-derived immortalized Schwann cells (IWARS1) are available from the corresponding author, Kazunori Sango.

#### Data and code availability

iTRAQ quantitative proteomics data have been deposited in the ProteomeXchange Consortium (http://proteomecentral.proteomexchange.org/cgi/GetDataset) via the jPOST partner repository (https://jpostdb.org/) with the dataset identifier PXD036003. The other datasets generated in this study are available from the corresponding author upon reasonable request.

### EXPERIMENTAL MODEL AND SUBJECT DETAILS

#### Drosophila melanogaster

All experimental subjects were female *Drosophila melanogaster*, except for males in Figure S3. The flies’ age was 14 days posteclosion if it is not stated in the figure legend or method details. Flies for strain maintenance were grown on a standard cornmeal-yeast-glucose diet (NSD in Table S1) on a 12:12-h light:dark cycle at 25°C and 60%–70% humidity. The fly lines were obtained from the Bloomington *Drosophila* Stock Center (BDSC) or the Vienna *Drosophila* Resource Center (VDRC).

#### Spontaneously immortalized mouse Schwann cells

Adult WT C57/BL6J mouse-derived immortalized Schwann cells (immortalized WT AR Schwann cells 1, IWARS1) (Niimi *et al*., 2021) were established as described previously (Niimi et al., 2018). The cells were maintained in Dulbecco’s modified Eagle’s medium (DMEM; 1 g/L glucose with L-Gln and Sodium Pyruvate, Nakalai Tesque Cat# 08456) containing 5% fetal bovine serum (FBS; Thermo Fisher Scientific Cat# 12483020) and 12.5 ng/ml Heregulin-β3, epidermal growth factor domain (Merck, Sigma-Aldrich Cat# 01-201). For ixazomib treatment, approximately 2 × 10^7^ cells were seeded on 100-mm plastic dishes and maintained in 5% FBS/DMEM for 48 h. The medium was then replaced with fresh 5% FBS/DMEM containing 10 ng/ml ixazomib citrate (MLN9708, Selleck Chemicals Cat# S2181) or 0.1% DMSO, and the cells were incubated for 24 h, rinsed with prewarmed phosphate-buffered saline (PBS) three times, and stored at –80°C.

## METHOD DETAILS

### Fly food recipes

The experimental HSD consisted of the NSD plus additional final concentrations of sucrose (Nacalai Tesque Cat# 30403-84) by volume (Table S1). HSD with 30% sucrose was used when the concentration is not mentioned. For drug administration, Formula 4-24® Instant *Drosophila* Medium, Blue food (Carolina Biological Supply Cat# 173214) was used; 1.2 g of instant *Drosophila* medium was hydrated with 4 ml of distilled water (dH_2_O) with or without 30% sucrose. Pioglitazone (Combi-Blocks Cat# QA-7806), metformin (Fujifilm Wako Pure Chemicals Cat# 138-18661), ixazomib, or DMSO was added to dH_2_O or 30% sucrose before food hydration. Trehalose (Merck, Sigma-Aldrich Cat# T9531), maltose (Fujifilm Wako Pure Chemicals Cat# 130-0615), glucose (Nacalai Tesque Cat# 16805-64), or fructose (Nacalai Tesque Cat# 16315-42) was dissolved in dH_2_O, and each solution was added to the Instant Blue food.

### Heat avoidance test

The heat avoidance test system developed by Manev and Dimitrijevic (2004) was modified for our experiment. A plastic tube (3 mm in diameter) was coiled around the test tube (acryl tube 10 mm in diameter, 100 mm in length, and 1 mm in thickness) to make a heat band zone by pumping temperature-controlled hot water (Perista pump, ATTO Cat# AC-2120; heater controller, WARNER Cat# TC-324B). The open end of the test tube was plugged with cotton to prevent the flies from escaping. The temperature of the heat band zone inside the test tube was measured with a digital thermometer (ANRITSU Cat# HA-100K).

First, the climbing test without the heat barrier (without perfusion) was performed to confirm the locomotor function of the test flies. After the flies are tapped down to the bottom of the tube, they climb up the wall of the tube by negative geotaxis. Flies that did not climb to the top of the tube within 10 sec were excluded from the heat avoidance test. The avoidance test with the heat barrier at 25°C, 38°C, 40°C and 42°C was then performed to determine the avoidance rate (%), which was calculated as the ratio of the number of flies that avoided the heat barrier to the number of flies tested. Seven to nine flies (one fly per one tube) were used for each experiment. The avoidance rate was calculated from two trials for each replicate. The interval between the tests was 2 min. For experiments using flies expressing the temperature-sensitive mutant *shi*^*ts1*^, the flies were kept at 30°C for 10 min before the assay, which was performed at 30°C (restrictive temperature). For experiments using flies expressing the temperature-sensitive mutants *Prosβ6*^*1*^ or *Prosβ2*^*1*^, the flies were maintained at 25°C or 29°C for 14 days, and the heat avoidance test was performed at room temperature.

### Hot plate test of decapitated flies

Decapitation and the noxious heat response test were performed according to a previously published protocol (Ohashi and Sakai, 2018), with slight modifications. Female flies or *pain*^*EP*^ flies fed NSD or HSD for 14 days were used for the experiment. Headless flies were prepared by cutting the heads of CO_2_–anesthetized flies off with microscissors. The length of anesthesia was limited to 3 min. The headless flies were placed in plastic vials at 25°C for 30–60 min. After recovering from anesthesia, a single fly was placed at the center of a thermo plate (8 × 10 cm) of a block incubator (ASTEC Cat# Bl-516S). The number of jumps or tumbles during 30 sec was counted, and the average number for nine flies was determined for each replicate. Flies that did not maintain their normal standing posture and did not respond to gentle mechanical stimulation with a brush were excluded.

### Climbing assay

The climbing assay was performed as described previously (Suzuki et al., 2015), with slight modifications. 10–20 flies were used for each genotype. The mean score of three trials was calculated for each replicate.

### Histological analyses

Fluorescence observation of adult fly legs was performed according to a previously published protocol (Guan et al., 2018), with slight modifications. CO_2_-anesthetized flies were kept in PBS containing 0.5% Triton X-100 (0.5PBT) for 10 min. The anterior or posterior legs were cut from the thorax and fixed in 10% formaldehyde (Polysciences Cat# 04018) overnight (14–16 h) at 4°C. The legs were washed with 0.5PBT three times for 20 min and then incubated in 50% glycerol in PBS for more than 2 h. The legs were mounted on a glass slide with SlowFade Diamond Antifade Mountant (Thermo Fisher Scientific Cat# S36963). Confocal microscopic images were obtained by a TCS-SP8 confocal microscope (Leica RRID: SCR_018169): mCD8-RFP + cuticle, ex 514 nm/em 584–616; cuticle, ex 488 nm/em 549–579. Postimaging processing to obtain the pure mCD8-RFP signal was performed with ImageJ/FIJI (Schneider et al., 2012) (RRID: SCR_002285), as described previously (Guan et al., 2018). The cell numbers (tarsi 5) and cell volumes (tarsi 4–5) of the mCD8-RFP-labeled neurons were analyzed by using Leica Application Suite X (Leica RRID: SCR_013673) and Imaris software (Bitplane RRID: SCR_007370), respectively. The legs of 4–10 flies were used for each group.

For LDs staining, the abdominal fat body was dissected from 2- or 14-day-old adult flies and fixed in 10% formaldehyde for 30 min. The tissues were washed with PBS three times, then incubated with 1 μM LipiDye (Lipid Droplet Green, Funakoshi Cat# FDV-0027) for 30 min on ice. After incubation with 1 μg/ml DAPI (FUJIFILM Wako Cat# 340-07971) in PBS for 5 min, the tissues were mounted with 80% glycerol.

### Measurement of hemolymph glucose and trehalose

Flies were starved for 15 h by placing them in a vial containing 0.75% agar. Hemolymph was pooled from 20 to 30 adult flies to obtain 1 µl for assay. The flies were punctured in the thorax with a fine needle and placed into 0.5-ml tubes whose bottoms had been punctured with a 22-gauge needle. The tubes were set into 1.5-ml tubes and centrifuged at 2600 × *g* for 5 min at 4°C. One microliter of hemolymph was diluted with 7 µl of citric acid buffer, pH 5.6, and heated at 95°C for 5 min. The glucose concentration was measured with LabAssay™ Glucose (FUJIFILM Wako Shibayagi Cat# 638-50971) according to the manufacturer’s protocol. Trehalose in the hemolymph was determined after digesting it to glucose by incubation with Trehalase (Merck, Sigma-Aldrich Cat# T8778) at 37°C for 14–16 h. The digested hemolymph was neutralized by adding 500 mM Tris, pH 7.5, before glucose assay.

### Insulin sensitivity experiment

The flies were starved for 16 h. The thorax was dissected, a vertical incision was made in the midline, and the thorax was placed in a microtube containing 200 μl of Schneider’s *Drosophila* Medium (Merck, Sigma-Aldrich Cat# S0146) on ice. After the planned number of thoraxes was collected, the microtubes were incubated for 10 min at 25°C. Two hundred microliters of medium containing 2 μM human insulin (Merck, Sigma-Aldrich Cat# I9278) or medium only was added to the thoraxes for 10 min (the final concentration of insulin was 1 μM). After the insulin treatment, the medium was removed, and the thoraxes were washed with ice-cold PBS and homogenized in 100 μl of 2 × sodium dodecyl sulfate (SDS) buffer (FUJIFILM Wako Cat# 196-16142). The lysates were heated at 95°C for 10 min, then centrifuged at 18,000 × *g* for 5 min. The supernatant protein samples were subjected to immunoblotting analysis.

### Quantitative RT-PCR

Total RNA was isolated from the heads of 5-day-old female flies (for *Ilp5/2/3* and *Gapdh*) or the whole bodies of 2-day-old (for *PSMD9* and *RpL32*) or 14-day-old (for *HSPs* and *RpL32*) flies by using TRIzol (Thermo Fisher Scientific Cat# 15596018). cDNA was generated with a PrimeScript RT reagent kit (Takara Cat# RR047A). Quantitative real-time polymerase chain reaction (qRT-PCR) was performed with the KAPA SYBR Fast qPCR kit (KAPA Biosystems Cat# KK4608) on a CFX96 Real-Time PCR Detection System (Bio-Rad RRID: SCR_018064). The relative amounts of transcripts were calculated by the standard curve method by using CFX Manager 3.1 software (Bio-Rad RRID: SCR_017251). The sequences of the primers were as follows:

*Ilp5* forward, 5’-AGAGAACTTTGGACCCCGTGAA-3’

*Ilp5* reverse, 5’-TGAACCGAACTATCACTCAACAGTCT-3’

*Ilp2* forward, 5’-GAATCACGGGATTATACTCCTCG-3’

*Ilp2* reverse, 5’-ATGAGCAAGCCTTTGTCCTTCA-3’

*Ilp3* forward, 5’-GAGGCACCTTGGGCCTATTC-3’

*Ilp3* reverse, 5’-CATGTGGTGAGATTCGGAGCTA-3’

*Gapdh* forward, 5’-TAAATTCGACTCGACTCACGGT-3’

*Gapdh* reverse, 5’-CTCCACCACATACTCGGCTC-3’

*dPSMD9* forward, 5’-CAACAGAAACGGACAGATCCTG-3’

*dPSMD9* reverse, 5’-CGCTGTGATATTGGTTGAGCAG-3’

*RpL32* forward, 5’-AGCGCACCAAGCACTTCATCCGCCA-3’

*RpL32* reverse, 5’-GCGCACGTTGTGCACCAGGAACTTC-3’

*DNAJ1* forward, 5’-TAGATATTGGCCGAGGATGC-3’

*DNAJ1* reverse, 5’-TCAGTCTCCAACACGAAACG-3’

*Droj2* forward, 5’-ATACTTGGCGTGAAGCCTAATG-3’

*Droj2* reverse, 5’-CACTTCGTAAGCCTGCGAAAT-3’

*Hsp70A* forward, 5’-CAAGATGCATCAGCAGGGTG-3’

*Hsp70A* reverse, 5’-AACAAAAATTAGACTCTTTGGCCT-3’

*Hsp70Bb* forward, 5’-AATCGCAGAGGACATGAAGC-3’

*Hsp70Bb* reverse, 5’-TCATCTTGGTCAGCACCATC-3’

*Hsp83* forward, 5’-TGTCATCCAGCACTTGAAGG-3’

*Hsp83* reverse, 5’-TGGACTTCATCAGCTTGCAC-3’

*Hsc70-1* forward, 5’-TTTCTGTGCTGACCATCGAG-3’

*Hsc70-1* reverse, 5’-TGGCCCAGATCCTTCTTATG-3’

### Body weight measurement

15–20 flies were put into preweighed microtubes. The total weight of the microtube was measured, and the body weight per fly was calculated.

### Lifespan Analysis

Experimental flies were raised with NSD, allowed to mate for 48 h after emerging, and then sorted with the use of CO_2_ anesthesia. The sorted flies were placed in vials containing the indicated media. The vials were changed without anesthesia to fresh media every 2– 3 days, and the number of dead flies was counted until all flies were dead.

### iTRAQ quantitative proteomics

Quantitative proteomic analyses of mouse Schwann cells were performed using isobaric tags for relative and absolute quantitation-mass spectrometry (iTRAQ-MS)-based platforms. For the preparation of protein extract, the cells were suspended in 8 M urea for 10 min on ice, transferred to plastic tubes, and sonicated (10 sec sonication/10 sec interval, repeated three times; Handy Sonic TOMY SEIKO Cat# UR-20P). The supernatant was cleared by centrifugation at 18,000 × *g* for 10 min at 4°C, and an equal amount of distilled water was added to reduce the urea concentration. Proteins were concentrated by using the Amicon Ultra centrifugal device (NMWL: 3K) (Merck, Millipore Cat# UFC500324), and the concentration was determined by the Pierce BCA protein assay kit (Thermo Fisher Scientific Cat# 23225). An equivalent mixture of three DMSO and four ixazomib samples was prepared for the reference sample. Fifty micrograms of protein from each sample were used for iTRAQ labeling according to the manufacturer’s instructions (iTRAQ Reagents – 8plex, SCIEX Cat# 4390811) and then fractionated into five fractions by cation exchange (Cation Exchange Buffer Pack, SCIEX Cat# 4326747). Each collected fraction was analyzed by the TripleTOF 5600^+^ system (SCIEX RRID: SCR_018053) equipped with the EASY-nLC 1200 system (Thermo Fisher Scientific Cat# LC140). The data were processed with ProteinPilot (ver. 5.0.1, SCIEX RRID: SCR_018681) against the UniProt 2010.06.22 database supplemented with 245 frequently observed contaminants, including human keratins, bovine serum proteins, and proteases. The Paragon algorithm (ver. 5.0.1.0, 4874) was used, with the following settings: a) iTRAQ 8plex (Peptide Labeled), b) methyl methanethiosulfonation of the cysteine residues, c) trypsin as enzyme, d) urea denaturation as special factors, e) *Mus musculus* as taxonomy. Default settings for iTRAQ isotope quantification, bias correction, and background correction were applied.

We identified 2190 proteins using a detection protein threshold (unused [Conf] cutoff) of > 1.3 (95%). The fold differences of identified proteins were determined as the iTRAQ label intensity ratios of the reference samples, and the data were exported to Excel files. The fold difference values were transformed into Log2. The values were normalized by quantile normalization using the preprocessCore R-package (Bolstad, 2021). Empirical Bayes moderated *t* tests were used to identify differentially expressed proteins in the ixazomib groups relative to the DMSO groups by using the limma R-package (Ritchie et al., 2015). Protein groups displaying Benjamini-Hochberg adjusted *P*-values (FDR q value) < 0.05 were considered differentially abundant. The heat map with hierarchical clustering was created by using the dendextend R-package with the ward D2 method (Galili, 2015). Enriched ontology clusters of differentially expressed proteins were identified by using Metascape (RRID: SCR_016620) (Zhou et al., 2019).

### Immunoblotting

Lysates of Schwann cells were prepared as described for the iTRAQ proteomics and mixed with equal volumes of 4× SDS buffer (FUJIFILM Wako Cat# 196-16142). The samples were separated by polyacrylamide gels, transferred to polyvinylidene fluoride (PVDF) membranes (Merck, Millipore Cat# IPVH00010), and blocked with 3% skim milk. Antibodies were diluted in Can Get Signal (TOYOBO Cat# NKB-201) immunoreaction enhancer solution. The signals were visualized with ImmunoStar Zeta or ImmunoStar LD (FUJIFILM Wako Pure Chemicals Cat# 297-72403, 297-69903), and images were captured by a bioimaging analyzer, LAS4000 (Fujifilm Life Science). Signal intensities were quantified by densitometry using ImageJ software. The antibodies used were as follows: anti-phospho-*Drosophila* Akt (Ser505) (Cell Signaling Technology Cat# 4054, 1:2000); anti-Akt (pan) (C67E7) (Cell Signaling Technology Cat# 4691, 1:2000); anti-Actin (Developmental Studies Hybridoma Bank Cat# jla20, 1:2000); anti-Multi Ubiquitin, FK2 (MBL International Cat# D058-3, 1:1000); anti-DNAJA1 (Merck, Sigma-Aldrich Cat# HPA001306, 1:2000); anti-HSP70, Clone C92F3-5 (StressMarq Biosciences Cat# SMC-100, 1:2000); anti-HSP90, Clone 16F1 (Novus Biologicals Cat# NBP1-97506, 1:1000); anti-PSMD9 (Bioworld Technology Cat# BS7754, 1:2000); horseradish peroxidase (HRP)-conjugated anti-rabbit IgG (Jackson ImmunoResearch Laboratories Cat# 111-035-003, 1:10000); HRP-conjugated goat anti-mouse IgG (Jackson ImmunoResearch Laboratories Cat# 115-035-003, 1:10000); and HRP-conjugated anti-mouse kappa light chain (Novus Biologicals Cat# NB7549, 1:10000).

## QUANTIFICATION AND STATISTICAL ANALYSIS

GraphPad Prism software (version 8.4.3, GraphPad Software RRID: SCR_002798) was used for statistical analyses and generation of graphs. The results are presented as means ± SEM. The types of statistical analyses, values of *N* (replicates), and animal numbers per *N* can be found in the Details of Methods and Figure Legends.

## SUPPLEMENTAL INFORMATION TITLES

**Document S1**. Figures S1–S5. Table S1–2.

**Table S3 related to Figure 3**. iTRAQ proteomics results. (A) All data for the proteins identified are shown. The values of fold difference of identified proteins were determined by the iTRAQ label intensity ratio of the reference sample. (B) List for the differentially expressed proteins with corresponding GO terms. Sixty-six proteins displaying Benjamini-Hochberg adjusted *P* value (false discovery rate [FDR] q value) of less than 0.05 were considered differentially abundant. (C) Protein list for significantly enriched GO Biological Processes terms. GO analysis was performed on the differentially expressed proteins using Metascape (Zhou et al., 2019). The GO terms with FDR q value < 0.05 was represented.

