## Supplementary material for "A *Drosophila* model of diabetic neuropathy reveals a crucial role of proteasome activity in the glia": Document S1

**Figure S1**

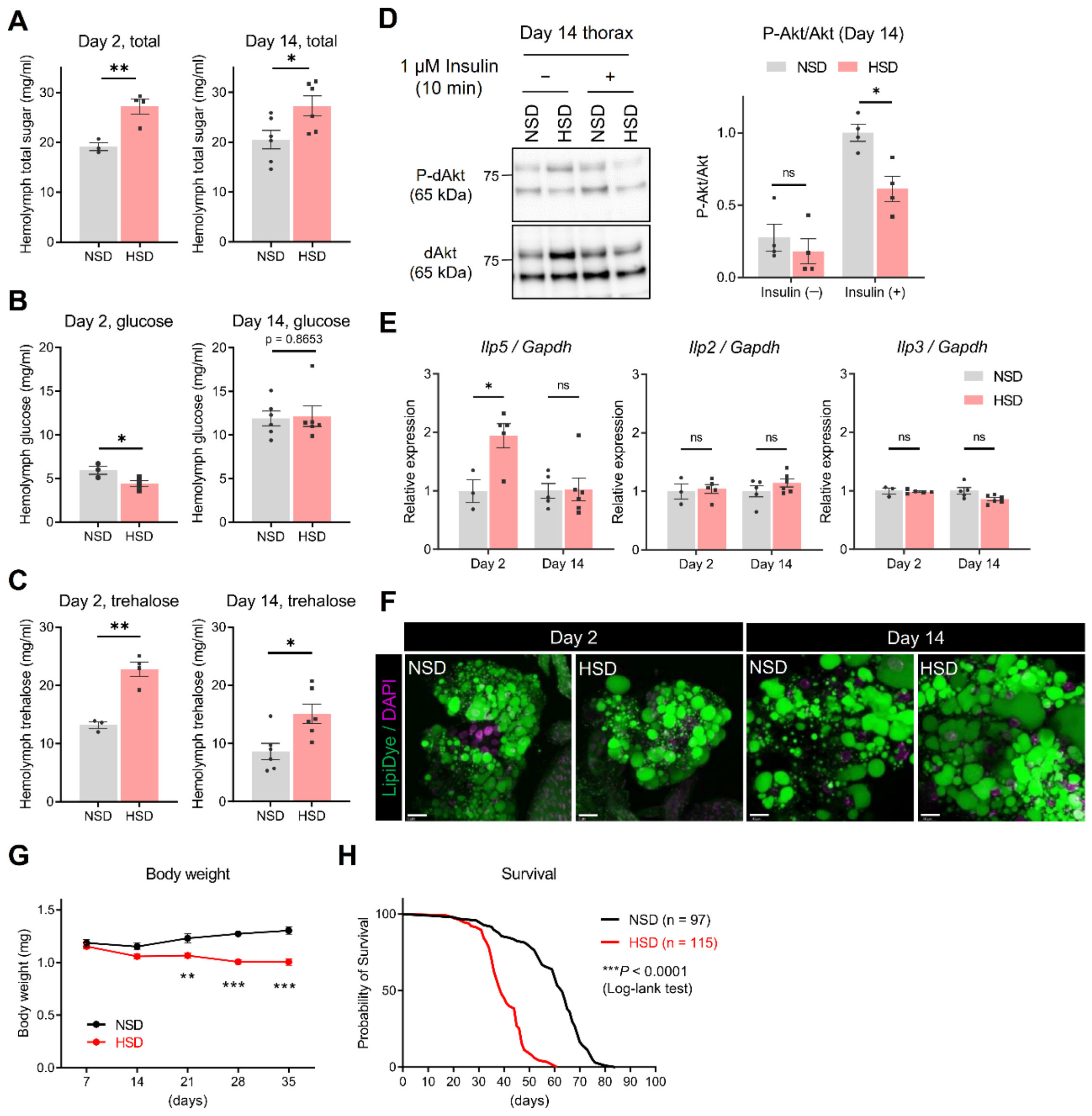


**Figure S1 related to Figure 1. A high-sugar diet induced diabetes-like metabolic features in wild-type female flies**

Wild-type (WT) flies were reared on a standard cornmeal-yeast-glucose diet (normal-sugar diet, NSD), and after the eclosion, female flies were collected and divided into the NSD group and HSD (high-sugar diet, NSD with an additional 30% sucrose) group. (A–C) HSD-induced hyperglycemia. Hemolymph total sugar (glucose + trehalose) (A), glucose (B) and trehalose (C) at day 2 and 14 of HSD feeding. Unpaired *t* test (N = 3–4 for day 2, N = 6 for day 14, 19–25 flies per replicate). (D) Insulin-induced *Drosophila* Akt (dAkt) phosphorylation was reduced by HSD. Thoraxes were dissected from starved flies at 14 days and treated either with insulin or vehicle in Schneider's *Drosophila* medium. Signal intensity was quantified from western blotting for phospho-*Drosophila*

Akt (P-dAkt) and total dAkt. Two-way ANOVA post hoc Tukey's test (N = 3–4 experiments). (E) Transcript levels of *Insulin-like peptide* (*Ilp*) 5, *Ilp2*, and *Ilp3* in flies at 2 and 14 days. Two-way ANOVA post hoc Tukey's test (N = 3–6, five heads per replicate). (F) HSD causes enlargement of lipid droplets (LDs) in the abdominal fat body. LDs were stained with LipiDye. (G) HSD gradually decreased body weight. Two-way ANOVA post hoc Tukey's test (N = 5, means of 4–20 flies were calculated for each replicate). (H) HSD significantly shortened lifespan. Log-lank test ($p < 0.0001$, 97 flies for NSD, 115 flies for HSD). ns, not significant, $*p < 0.05$, $**p < 0.01$, $***p < 0.001$.

Figure S2

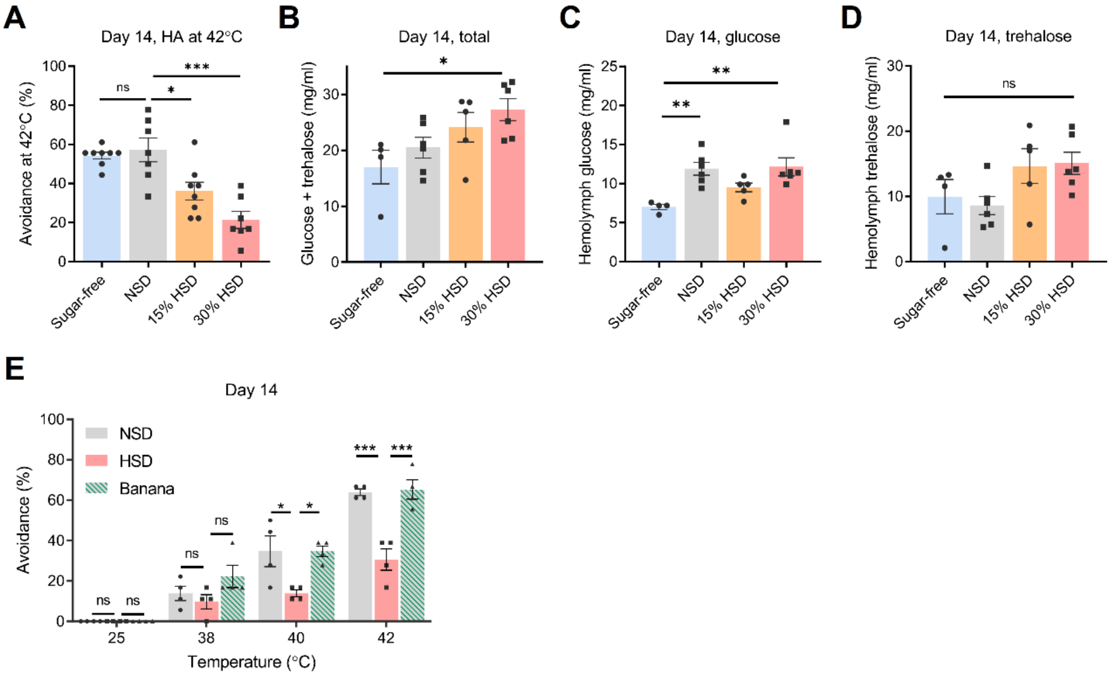


**Figure S2 related to Figure 1. Dose-dependent effects of dietary carbohydrate on heat avoidance behavior and hemolymph sugar**

(A–D) Female flies were fed indicated diets for 14 days, and heat avoidance at 42°C (A) and hemolymph sugar concentrations (B–D) were analyzed. Sugar-free, NSD without glucose; 15 or 30% HSD, NSD with 15 or 30 % sucrose. One-way ANOVA post hoc Tukey's test (N = 4–8, 7–9 flies per replicate for A, 19–25 flies per replicate for B–D). (E) Flies fed banana showed similar heat avoidance profiles to flies fed NSD. Two-way ANOVA post hoc Tukey's test (N = 4, 7–9 flies per replicate). ns, not significant, *$p < 0.05$, **$p < 0.01$, ***$p < 0.001$.

### Figure S3

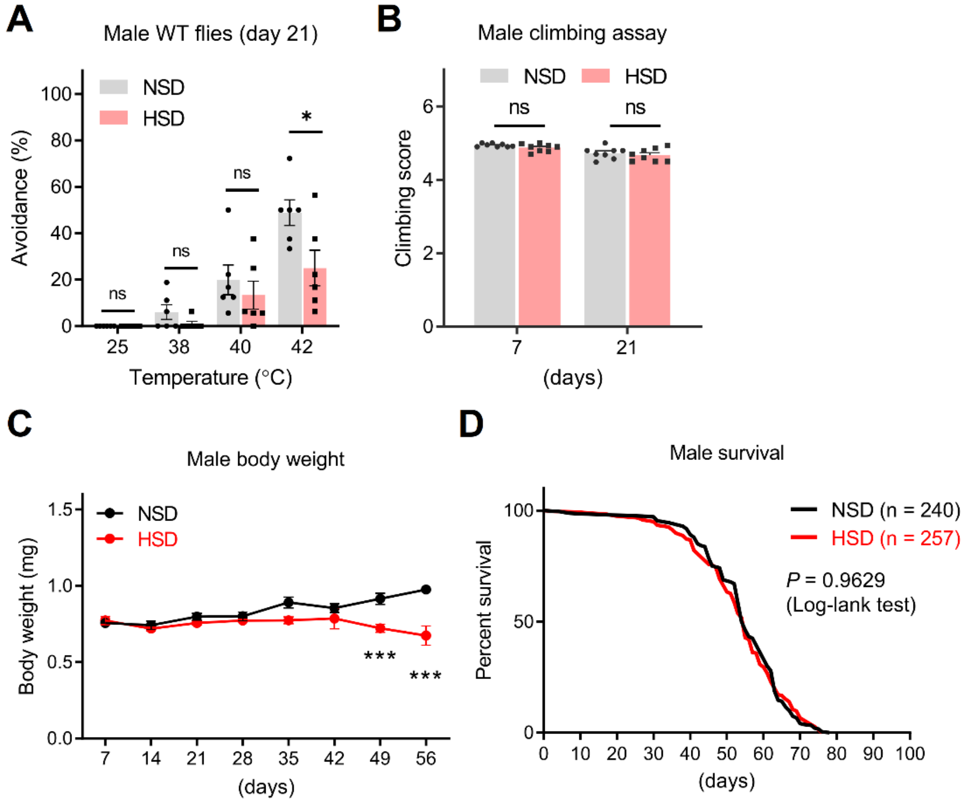


**Figure S3 related to Figure 1. Effects of HSD on male WT flies**

(A) Heat avoidance test of male WT flies at day 21. Two-way ANOVA post hoc Tukey's test (N = 6, 7–9 flies per replicate). (B) Climbing assay. Two-way ANOVA post hoc Tukey's test (N = 8, 10–20 flies per replicate). (C) Body weight. Two-way ANOVA post hoc Tukey's test (N = 3–5, means of 7–20 flies were calculated for each replicate). (D) Survival curve. Log-lank test (p = 0.9629, 240 flies for NSD, 257 flies for HSD). ns, not significant, *$p < 0.05$, ***$p < 0.001$.

### Figure S4

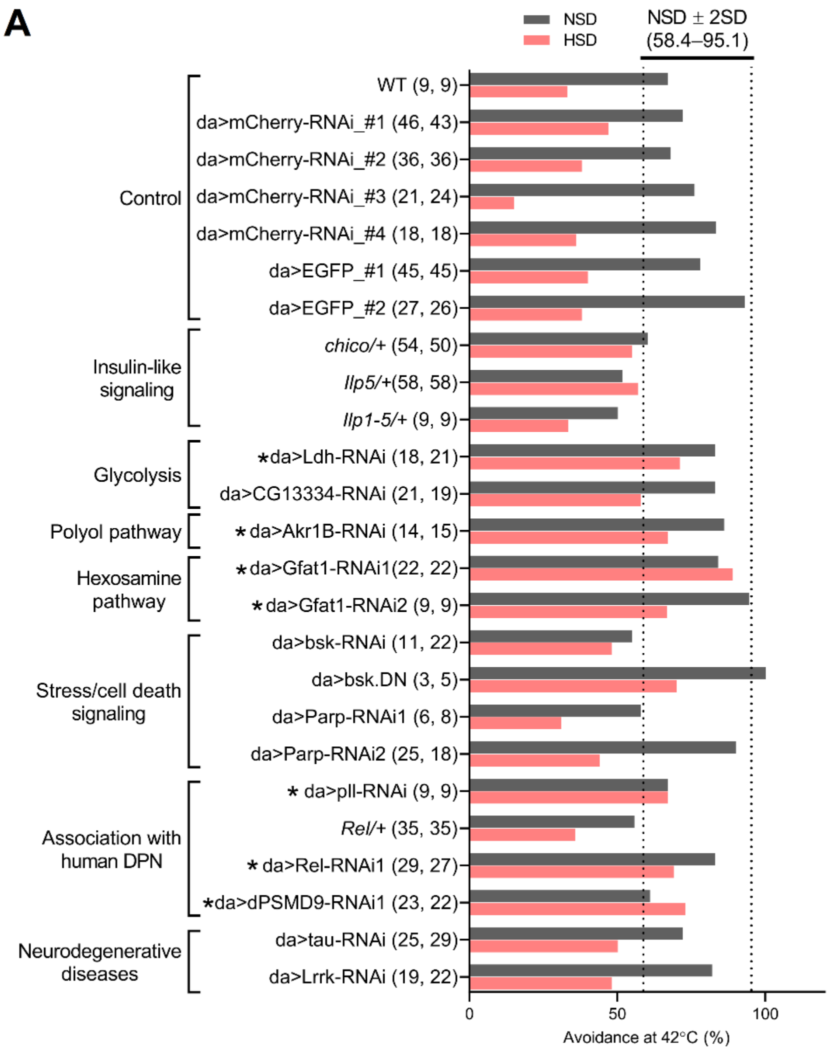


**B**

| Fly line | % HA (NSD) | % HA (HSD) | Difference (NSD−HSD) |
|---|---|---|---|
| da>dPSMD9-RNAi1 | 61.0 | 73.0 | -12.0 |
| da>Gfat1-RNAi1 | 84.0 | 89.0 | -5.0 |
| da>pll-RNAi | 67.0 | 67.0 | 0.0 |
| da>Ldh-RNAi | 83.0 | 71.0 | 12.0 |
| da>Rel-RNAi1 | 83.0 | 69.0 | 14.0 |
| da>Akr1B-RNAi | 86.0 | 67.0 | 19.0 |
| da>Gfat1-RNAi2 | 94.4 | 66.7 | 27.7 |

**Figure S4 related to Figure 3. Candidate genetic screening for modifiers of HSD-induced heat avoidance impairment**

(A) Screening for genetic modifiers of HSD-induced heat avoidance impairment. RNAi-mediated knockdown or overexpression flies under the control of the *daughterless* (*da*)-*GAL4*, or heterozygote mutant flies were raised on NSD or HSD for 14 days after eclosion and used for the heat avoidance test. *da-GAL4>mCherry-RNAi* (da>mCherry-RNAi #1–4) and *da-GAL4>EGFP* (da>EGFP #1–2) flies were used as control. The numbers in parentheses indicate the fly numbers used for NSD or HSD, respectively. The reference range was set as 58.4%–95.1 %, and seven fly lines, both of which heat avoidance rates for NSD and HSD are included in the reference range, were identified as hits (asterisks). (B) Hit fly lines were listed in order of the decreasing means difference between NSD and HSD.

### Figure S5

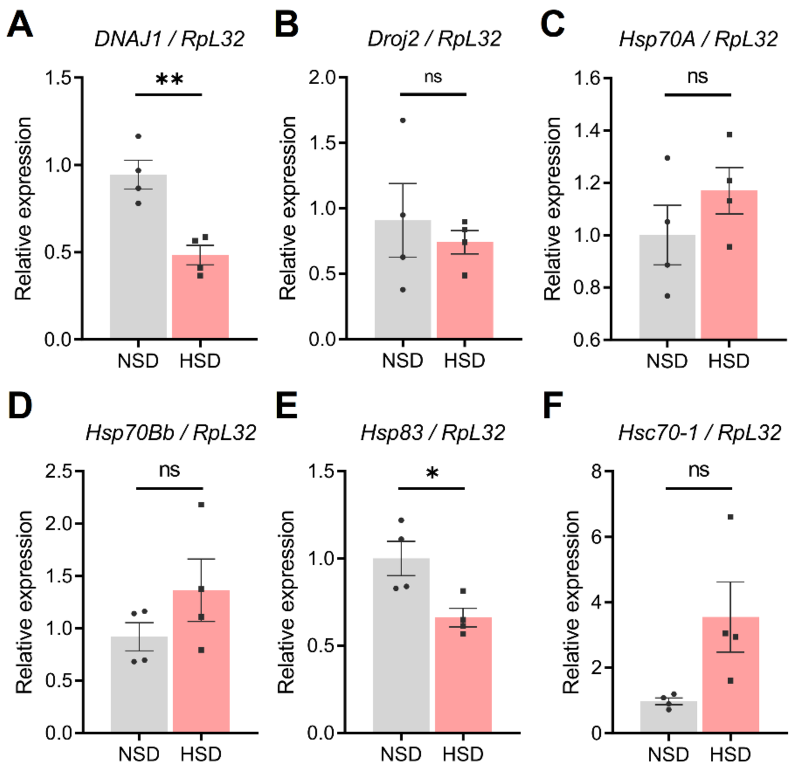


**Figure S5 related to Figure 6. Expression levels of HSPs in the HSD-fed flies**

(A–F) Expression levels of HSPs in the HSD-fed flies. Total RNA was isolated from whole bodies of 14-days-old flies. *RpL32* was used as an internal control. Student *t* test for (A) *DNAJ1*, (B) *Droj2*, (C) *Hsp70A*, (D) *Hsp70Bb*, and (E) *Hsp83*. Welch's *t* test for (F) *Hsc70-1*. N = 4, two flies per replicate.

### Table S1 related to STAR METHODS. *Drosophila* diet recipes

**Cornmeal-yeast-glucose diets (per L of media)**

| | NSD | HSD (30%) | HSD (15%) | Sugar-free | Banana |
|---|---|---|---|---|---|
| Sucrose (Nacalai Tesque Cat# 30403-84) | – | 300 g | 150 g | – | |
| Glucose (Nacalai Tesque Cat# 16805-64) | 100 g | | | – | |
| Agar (Ina Food Industry Cat# BA-10) | 7.5 g | | | | |
| Baker's yeast (Oriental Yeast) | 35 g | | | | |
| Cornmeal (Oriental Yeast) | 55 g | | | | |
| Bokinin* (Nacalai Tesque Cat# 06327-15) | 10 mL | | | | |
| Propionic acid (FUJIFILM Wako Cat# 163-04726) | 5 mL | | | | |
| $H_2O$ | up to 1 L | | | | |
| Total kcal | 737 | 1933 | 1335 | 377 | 930 |
| % Carbohydrate (w/v) | 15.5 | 45.4 | 30.4 | 5.5 | 22.5 |
| % Fat (w/v) | 0.5 | | | | 0.2 |
| % Protein (w/v) | 1.8 | | | | 1.1 |

*10 % (w/v) butyl 4-hydroxybenzoate in ethanol.

**Instant blue food (per φ24×H95mm vial)**

| | NSD | HSD |
|---|---|---|
| Formula 4-24® Instant *Drosophila* Medium, Blue food (Carolina Biological Supply Cat# 173214) | 1.2 g | |
| $H_2O$ | 4 mL | – |
| 30% sucrose in $H_2O$ | – | 4 mL |

**For saccharides experiment (Fig. 1D), the indicated saccharide solution was used instead of 30% sucrose solution.

***Vehicles (DMSO and ethanol), pioglitazone, metformin, ixazomib, and RU486 stock solutions were added to $H_2O$ or 30% sucrose in $H_2O$.

**Table S2 related to Figure 3. Candidate genes for genetic screening**

Candidate genes were selected by functions in glucose metabolism (insulin-like signaling, glycolysis, polyol pathway, and hexosamine pathway), stress and cell death signaling, association with human DPN, and neurodegenerative diseases. Asterisks indicate human genes that have been reported to be associated with DPN.

| Features | Fly gene | Human orthologs | Known functions, related reference(s) |
|---|---|---|---|
| Glucose metabolism | *chico* | *IRS1* | A substrate of the Insulin-like receptor; control of cell size and growth; related to HSD-induced cardiomyopathy in *Drosophila* (Na et al., 2013) |
| | *Insulin-like peptide 5* (*Ilp5*) | *Insulin,* *Insulin-like growth factor* | A peptide involved in the insulin signaling pathway |
| | *Lactate dehydrogenase* (*Ldh*) | *LDHA,* *LDHB* | Involved in glycolysis, gluconeogenesis, and pyruvate metabolism |
| | *CG13334* | *LDHAL6A/B/C* | Predicted to enable L-lactate dehydrogenase activity |
| | *Aldo-keto reductase 1B* (*Akr1B*) | *AKR1B1*,* *AKR1B10,* *AKR1B15,* *AKR1D1,* *AKR1C4,* *AKR1C2* | Predicted to aldose reductase, indanol dehydrogenase, and D-threo-aldose 1-dehydrogenase activity; Human ortholog *AKRB1* implicated in several diseases including DPN (Heesom et al., 1998; Sivenius et al., 2004; Gupta et al., 2017) |
| | *Glutamine-fructose-6-phosphate aminotransferase 1* (*Gfat1*) | *GFAT2,* *GFAT1* | Predicted to enable carbohydrate derivative binding activity and glutamine-fructose-6-phosphate transaminase activity; related to HSD-induced cardiomyopathy and podocyte dysfunction in *Drosophila* (Na et al., 2013, 2015) |
| Stress and cell death signaling | *basket (bsk)* | *MAPK8,* *MAPK10* | Encodes a serine/threonine-protein kinase, a key component of JNK pathway; regulation of stress response |
| | *Poly-(ADP-ribose) polymerase (Parp)* | *PARP1** | Encodes a nuclear enzyme modifying acceptor proteins through assembly of poly(ADP-ribose) polymers; Human ortholog is implicated in DPN (Nikitin et al., 2008) |
| Association with human DPN | *Pelle (pll)* | *IRAK4* | Encodes a serine-threonine protein kinase that functions in the Toll pathway. Human Toll-like receptor 4 is implicated in DPN (Rudofsky et al., 2004) |
| | *Relish (Rel)* | *NFKB1* | Encodes a transcription factor and the downstream component of the immune deficiency pathway, which regulates the antibacterial response. Human ortholog mediate a signal from toll-like receptor |
| | *CG9588* (*PSMD9*) | *PSMD9** | A chaperone during the assembly of the 26S proteasome. Human ortholog is implicated in DPN (Gragnoli, 2011; Salcini et al., 2021) |
| Neurodegenerative diseases | *tau* | *MAPT* | Microtubule-associated protein; human ortholog protein is related to several neurodegenerative diseases such as Alzheimer's disease, Pick's disease, frontotemporal dementia, cortico-basal degeneration, and progressive supranuclear palsy |
| | *Leucine-rich repeat kinase* (*Lrrk*) | *LRRK1,* *LRRK2* | Encodes a large Ser/Thr kinase involved in mRNA translational control, human ortholog protein LRRK2 is related to Parkinson's disease |
